# Darapladib, an inhibitor of Lp-PLA2, sensitizes cancer cells to ferroptosis by remodeling lipid metabolism

**DOI:** 10.1101/2023.04.08.536136

**Authors:** Mihee Oh, Seo Young Jang, Ji-Yoon Lee, Jong Woo Kim, Youngae Jung, Jinho Seo, Tae-Su Han, Eunji Jang, Hye Young Son, Dain Kim, Min Wook Kim, Kwon-Ho Song, Kyoung-Jin Oh, Won Kon Kim, Kwang-Hee Bae, Yong-Min Huh, Baek-Soo Han, Sang Chul Lee, Geum-Sook Hwang, Eun-Woo Lee

## Abstract

Arachidonic and adrenic acids in the membrane play key roles in ferroptosis, but how these fatty acids are manipulated in cells is largely unknown. Here, we reveal that lipoprotein-associated phospholipase A2 (Lp-PLA2) controls intracellular phospholipid metabolism and contributes to ferroptosis resistance. A metabolic drug screen identified that darapladib (SB-480848), an inhibitor of Lp-PLA2, synergistically induced ferroptosis with GPX4 inhibitors. Notably, darapladib was able to enhance ferroptosis under lipoprotein-deficient or serum-free conditions. Furthermore, Lp-PLA2 was located in the membrane and cytoplasm and suppressed ferroptosis, suggesting the critical role of intracellular Lp-PLA2. Lipidomic analysis showed that phosphatidylethanolamine (PE) species were generally enriched, while lysophosphatidylethanolamine (lysoPE) and free fatty acid levels were reduced, upon darapladib treatment. Finally, combination treatment with darapladib and PACMA31, a GPX4 inhibitor, efficiently inhibited tumor growth in a xenograft model. Our study suggests that inhibition of Lp-PLA2 is a potential therapeutic strategy to enhance ferroptosis in cancer treatment.

## Introduction

Ferroptosis is a newly identified form of programmed necrosis that requires free active iron (Dixon et al., 2012; Hassannia et al., 2019). Excessive accumulation of reactive oxygen species (ROS) in membrane phospholipids (PLs) is a main cause of ferroptosis; thus, lipid ROS are a distinct hallmark of ferroptosis (Aldrovandi et al., 2021). Lipid ROS are primarily controlled by glutathione peroxidase 4 (GPX4), an enzyme that directly reduces lipid peroxides in membrane phospholipids to lipid alcohols (Ingold et al., 2018; Seiler et al., 2008). Glutathione (GSH) is simultaneously oxidized by GPX4, thus acting as an essential cofactor of GPX4 (Lee et al., 2021b). Intracellular levels of GSH, which consists of cysteine, glutamate, and glycine, are mainly controlled by the cystine–glutamate antiporter. Based on this, several ferroptosis-inducing agents (FINs) that target GPX4 or cystine-glutamate antiporter have been developed and are regarded as potential drugs for cancer treatment (Hassannia *et al*., 2019; Lei et al., 2022). While the GPX4/GSH system protects most types of cells from ferroptosis, recent studies suggest that chemoresistant cancer cells with mesenchymal characteristics or drug-tolerant cancer cells are particularly susceptible to ferroptosis (Hangauer et al., 2017; Lee et al., 2020; Rodriguez et al., 2022; Viswanathan et al., 2017). Therefore, ferroptosis has been widely recognized as an emerging anticancer treatment strategy. In addition, ferroptosis is also implicated in various human diseases such as ischemia-reperfusion injury in the kidneys, heart, brain, and liver; thus, ferroptosis inhibitors have been developed for the treatment of these diseases (Jiang et al., 2021a).

As ferroptosis is induced by lipid peroxidation, various metabolic and signaling pathways that affect lipid peroxidation, such as lipid metabolism, iron metabolism, and antioxidant pathways, are involved in the ferroptosis pathway (Jiang *et al*., 2021a; Kim et al., 2021; Lee et al., 2021a; Zhang et al., 2022; Zheng and Conrad, 2020). In particular, arachidonic acid (AA; C20:4) and adrenic acid (AdA; C22:4), which are linked in membrane phospholipids, especially phosphatidylethanolamine (PE) and phosphatidylcholine (PC), are the most critical lipids for lipid peroxidation and ferroptosis (Doll et al., 2017a; Kagan et al., 2017). In this regard, the critical roles long-chain acyl-CoA synthetase 4 (ACSL4) and Lysophospholipid acyltransferase (LPCAT3), which mediate the incorporation of AA into PE or PC, in ferroptosis in various pathological contexts have been the focus of considerable research (Hassannia *et al*., 2019; Lee *et al*., 2021b; Tang and Kroemer, 2020). Under normal conditions, AA is synthesized from the n-6 essential fatty acid linoleic acid (LA; C18:2) in the liver and is supplied to tissues. In tumor, cancer cells also directly synthesize AA from LA, the most abundant polyunsaturated fatty acid (PUFA) in serum and plasma (Lee *et al*., 2020). AA can also be directly imported from extracellular fluid supplied from the liver via blood vessels and then further elongated into the AdA ELOVL5 (Aldrovandi *et al*., 2021; Lee *et al*., 2021b). While intracellular AA levels are important for ferroptosis, but only a small fraction of AA in phospholipids is oxidized under ferroptotic conditions (Doll *et al*., 2017a; Kagan *et al*., 2017). However, little is known about how cells maintain phospholipids containing AA to regulate lipid peroxidation and ferroptosis.

Phospholipase A2s (PLA2s) are enzymes that hydrolyze fatty acids at the sn-2 position of phospholipids, and there are more than 50 enzymes in the PLA2 superfamily (Peng et al., 2021). PLAs are largely classified into secretory PLA2 (sPLA2), cytosolic PLA (cPLA2), Ca^2+^-independent PLA2 (iPLA2), and lipoprotein-associated phospholipase A2 (Lp-PLA2; encoded by PLA2G7 and known as plasma platelet-activating factor acetylhydrolase [PAF-AH]), all of which have different sequences and structures (Burke and Dennis, 2009; Murakami et al., 2020). The major role of PLA2 is to extract AA from membrane phospholipids, thereby contributing to the formation of pathophysiological lipid mediators such as prostaglandins, leukotrienes, and lysophospholipids. Since AAs play a key role in ferroptosis, PLA2 is thought to be involved in ferroptosis. While cPLA2 is already known to increase lipid peroxide by releasing AA, its contribution to ferroptosis is unclear (Nanda et al., 2007; Wang et al., 2021a). Recently, iPLA2β, which is encoded by PLA2G6, was identified to specifically cleave oxidized PE containing AA (15-HpETE-PE), thereby suppressing ferroptosis (Beharier et al., 2020; Chen et al., 2021; Sun et al., 2021).

In this study, we discovered that darapladib, an Lp-PLA2 inhibitor, sensitizes cells to ferroptosis and shows antitumor activity when administered with PACMA31, a GPX4 inhibitor, in vivo. Using lipidomic analysis, we found that darapladib rewires lipid metabolism to render cells vulnerable to ferroptosis. Interestingly, intracellular Lp-PLA2, but not extracellular Lp-PLA2, seems to protect cells from ferroptosis. Thus, these results suggest that Lp-PLA2 is an essential negative regulator of ferroptosis sensitivity.

## Results

### A metabolic library screen identified darapladib, an Lp-PLA2 inhibitor, as a ferroptosis-sensitizing drug

To identify the metabolic pathways regulating ferroptosis and discover metabolic drugs that kill cancer cells synergistically with ferroptosis inducers, 403 metabolism-modulating compounds were screened from libraries on the basis of their ability to modulate RSL3-induced ferroptosis (Fig. 1a). In this screen, we used Hs746T cells, mesenchymal-type gastric cancer cells that are refractory to standard therapy but are sensitive to ferroptosis, as shown in our previous study (Lee *et al*., 2020). When used alone, several compounds reduced cell viability to less than 50%, but most compounds did not significantly alter the overall survival rate (Supplementary Fig. 1a). We focused on several compounds that have no toxicity when used alone but significantly promote cell death when administered with RSL3 and selected several candidates that acted synergistically with RSL3 (Supplementary Fig. 1a). After validation with ferrostatin-1 (Fer-1), a ferroptosis inhibitor, and investigation of the literature regarding known targets, mechanisms, in vivo use, and clinical trials, darapladib (SB-480848), an Lp-PLA2 inhibitor, was chosen as the final candidate (Supplementary Fig. 1b). Since darapladib itself is toxic to cancer cells when used at high concentrations, 2 μM darapladib was used in the following experiment (Supplementary Fig. 2a).

**Fig. 1.**
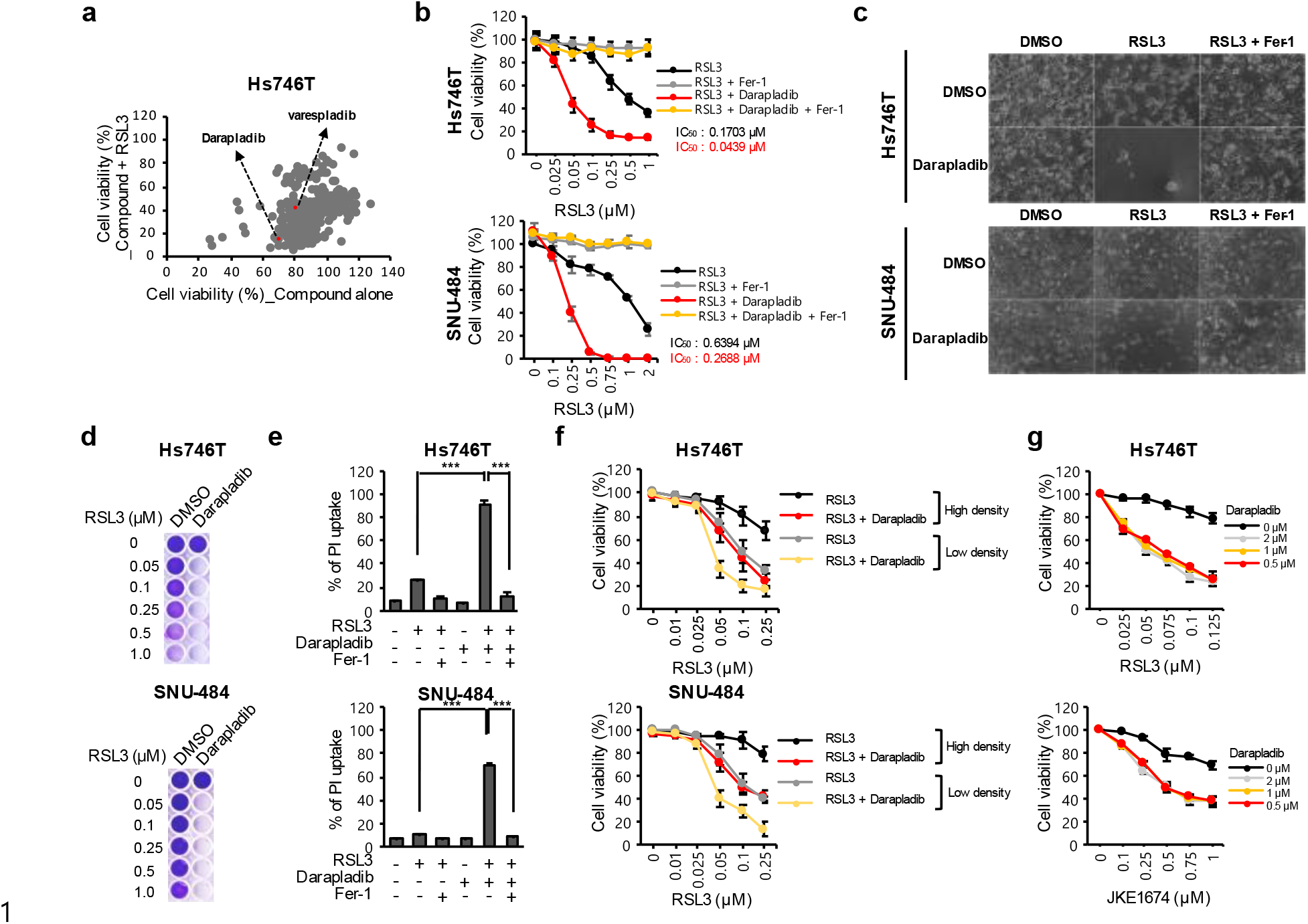
Metabolic library screening identified darapladib as a ferroptosis-targeting drug. (a) Relative viability of Hs746T cells treated with 10 μM compound alone or compound and 0.5 μM RSL3 for 24 h. (b) Relative viability of Hs746T and SNU-484 cells treated with increasing concentrations of RSL3 and/or 2 µM darapladib for 20 h. Cells were plated at 30,000 Hs746T cells/well and 40,000 SNU-484 cells/well in 200 μL media. (c) Images of cells treated with 0.2 μM RSL3 and/or 2 μM darapladib in the presence or absence of 1 μM Fer-1. (d) Crystal violet staining of cells treated with RSL3 and/or 2 μM darapladib for 48 h. Cells were plated at 20,000 Hs746T cells/well and 25,000 SNU-484 cells/well in 200 μL media. (d) PI uptake from Hs746T and SNU-484 cells treated with RSL3 and/or 2 μM darapladib in the presence or absence of 1 μM Fer-1 for 20 h. (e) Relative viability of cells at high (30,000 Hs746T cells/well and 40,000 SNU-484 cells/well) and low (20,000 Hs746T cells/well and 25,000 SNU-484 cells/well) density upon RSL3 and 2 μM darapladib treatment. (f) Relative viability of Hs746T cells treated with RSL3 or JKE1674 in the presence of increasing concentrations of darapladib. The data are the means ± SDs (n = 3 independent experiments).

Darapladib sensitized Hs746T and SNU-484 cells, both of which are mesenchymal-type gastric cancer cells, to ferroptosis (Fig. 1b-e) (Lee *et al*., 2020). Notably, when cells were cotreated with darapladib, the half-maximal inhibitory concentration (IC50) values for RSL3 were drastically lowered in both cell lines (Fig. 1b). The dramatic decrease in cell viability observed upon the combined treatment with RSL3 and darapladib was almost completely eliminated by treatment with Fer-1, a ferroptosis inhibitor (Fig 1b,c and Supplementary Fig. 2b). However, the pan-caspase inhibitor zVAD-fmk or the RIPK1 inhibitor necrostatin-1 (Nec-1) failed to rescue cell viability (Supplementary Fig. 2b). We confirmed that the decrease in cell viability was caused by necrotic cell death using PI uptake and LDH level assays (Fig. 1e and Supplementary Fig. 2c). In addition, RSL3 induced cell death more efficiently when cells were plated at a low density, as previously reported (Fig. 1f) (Vucetic et al., 2020). Under this condition, cell death was further enhanced by darapladib (Fig. 1f). Furthermore, 0.5 μM darapladib was sufficient to enhance GPX4 inhibitor-induced ferroptosis (Fig. 1g). These data suggest that darapladib specifically promotes ferroptosis induced by GPX4 inhibitors.

### Darapladib is a general activator of ferroptosis in various types of cells

When ferroptosis was induced with other GPX4 inhibitors, such as ML210 and JKE1674, a derivative of ML210, darapladib also enhanced ferroptosis in Hs746T and SNU-484 cells (Fig. 2a). Of note, the increase in cell death caused by the combined treatment was fully reversed by Fer-1, confirming that ferroptosis was indeed augmented by darapladib (Fig. 2b). In addition to gastric cancer cell lines, consistent synergistic effects of RSL3 and darapladib were observe in lung cancer cell lines such as A549 and H1299 and in the liver cancer cell line HepG2 (Supplementary Fig. 3a). We also found that ferroptotic cell death was promoted when RSL3 and darapladib were coadministered to YCC-16 cells, another type of anticancer treatment–refractory cell (Figure S3a). Furthermore, darapladib also renders non-cancer cells, such as H9c2 and MEFs, to RSL3-induced ferroptosis, suggesting the conserved effect of darapladib on ferroptosis (Supplementary Fig. 3b)

**Fig. 2.**
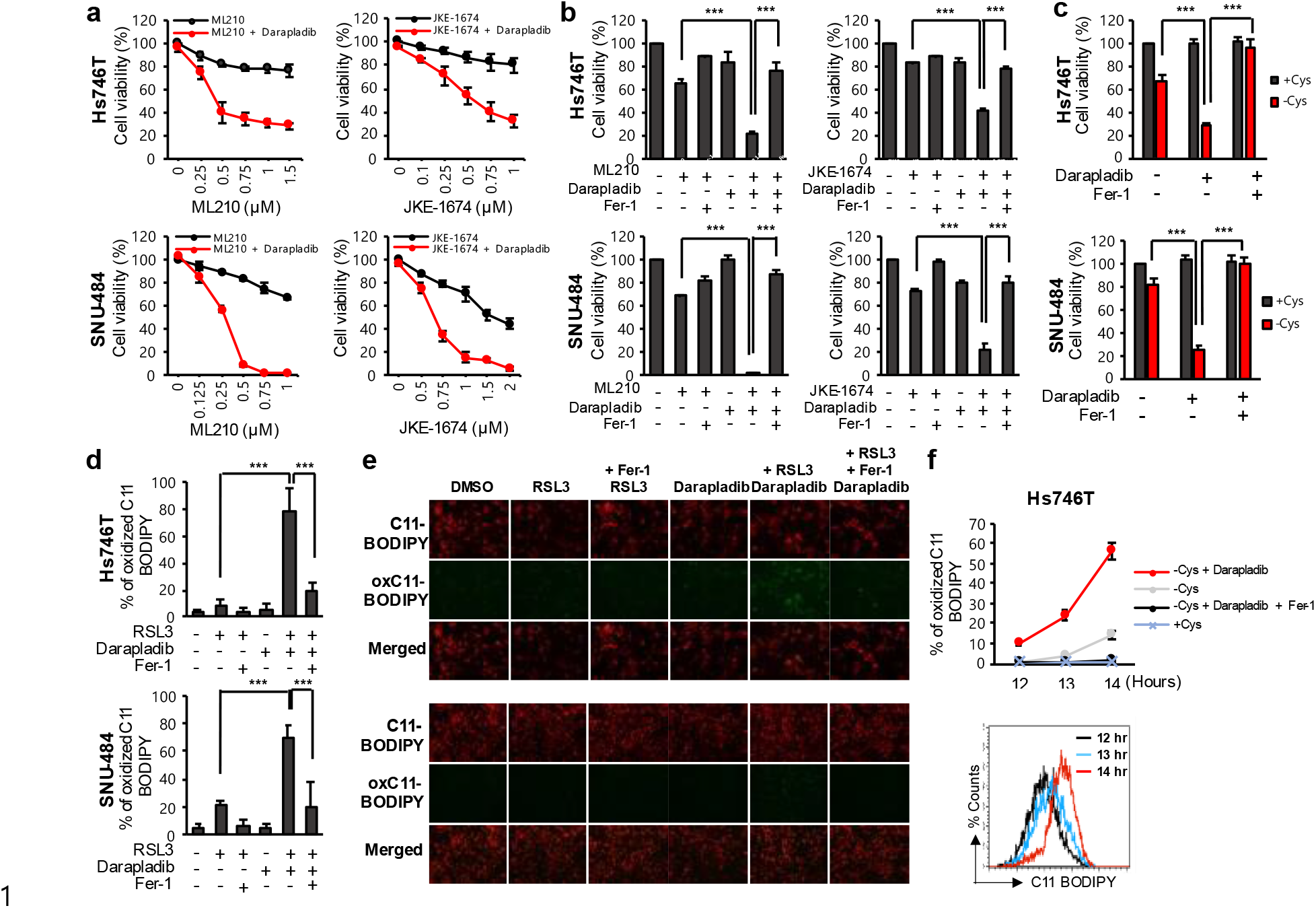
Darapladib sensitizes cells to ferroptosis induced by GPX4 inhibition or cysteine deprivation. (a) Relative viability of Hs746T and SNU-484 cells treated with increasing concentrations of ML210 or JKE1674 in the presence or absence of 2 μM darapladib for 20 h. (b) Relative viability of cells in the presence or absence of 1 μM Fer-1. (c) Cell viability of Hs746T and SNU-484 cells cultured in cysteine-deficient medium in the presence and absence of 2 μM darapladib for 18 h. (d and e) Lipid peroxidation levels in Hs746T and SNU-484 cells treated with 0.2 μM RSL3 and 2 μM darapladib for 1 h. Lipid oxidative potential was assessed by flow cytometry (d) and microscopy using C11-BODIPY^581/591^ (e). Probes fluorescing green represent those that have been oxidized. (f) Lipid peroxidation levels in Hs746T cells cultured in cysteine-deficient medium in the presence or absence of darapladib for 12∼14 h. The data are the means ± SDs (n = 3 independent experiments).

Next, we investigated whether darapladib also responds to other ferroptotic stimuli, such as GSH depletion. Culture in cysteine-depleted medium for 18 h did not greatly reduce the cell viability, but culture in cysteine-depleted medium in the presence of darapladib caused rapid cell death in both Hs746T and SNU-484 cells (Fig. 2c). Again, cell viability was remarkably recovered by Fer-1, suggesting that darapladib also sensitizes cells to cysteine deprivation-induced ferroptosis (Fig. 2c). In addition, darapladib can promote ferroptosis induced by erastin, an inhibitor of system x_c_^-^, (Supplementary Fig. 3c).

Next, we measured the level of lipid peroxidation, a hallmark of ferroptosis, using C11 BODIPY 581/591. When cells were treated with 0.1 uM RSL3, no increase in lipid peroxidation compared to that in the control group was detected. However, more than 80% of cells cotreated with darapladib and RSL3 showed oxidized C11-BODIPY, as evidenced by flow cytometry and fluorescence microscopy (Fig. 2d, e). These signals were completely abolished by Fer-1, confirming that lipid peroxidation was indeed enhanced. In addition, while oxidized C11-BODIPY signals transiently appeared before cells died at approximately 14 h after cysteine deprivation, darapladib facilitated lipid peroxidation, as we observed oxidized C11-BODIPY signals at approximately 12 h (Fig. 2f). These data suggest that darapladib renders cells sensitive to ferroptosis by increasing lipid peroxidation.

### Darapladib sensitizes cells to ferroptosis independent of lipoprotein

We next investigated whether darapladib sensitizes cells to ferroptosis in an Lp-PLA2-dependent manner. Since Lp-PLA2 is known to be associated with lipoprotein (Tellis and Tselepis, 2009), we first tested the effect of darapladib in lipoprotein-deficient conditions. When cells were cultured in lipoprotein-deficient human serum (LPDS), RSL3-induced ferroptosis was drastically diminished, and darapladib had no further effect when administered for 6 h (Fig. 3a, b). While starvation stress may activate mTOR pathway that can suppress ferroptosis, the levels of phospho-S6K were unaffected by lipoprotein deficiency (Supplementary Fig. 4a) (Yi et al., 2020; Zhang et al., 2021b). Interestingly, supplementation with HDL, but not with LDL or VLDL, re-sensitizes cells to ferroptosis under lipoprotein deficiency, suggesting that HDL may contribute to ferroptosis although underlying mechanism is unclear and further lipidomic analysis will be required (Supplementary Fig. 4b). Notably, cells cultured in LDPS for 24 h with darapladib for 24 h died when treated with RSL3, while control cells did not (Fig. 3c). These data imply that although lipoprotein deficiency generally slows the ferroptosis response, darapladib is still able to increase ferroptosis in the absence of lipoprotein. In addition, darapladib sensitized cells to ferroptosis under serum starvation conditions, supporting the indispensable role of lipoprotein in darapladib-mediated ferroptosis sensitization (Fig. 3d). Interestingly, unlike lipoprotein deficiency, serum starvation did not alleviate ferroptosis, probably due to the concomitant depletion of anti-ferroptotic components in the serum such as vitamin E, selenium, and CoQ10 or a difference in composition between human and bovine serum.

**Fig. 3.**
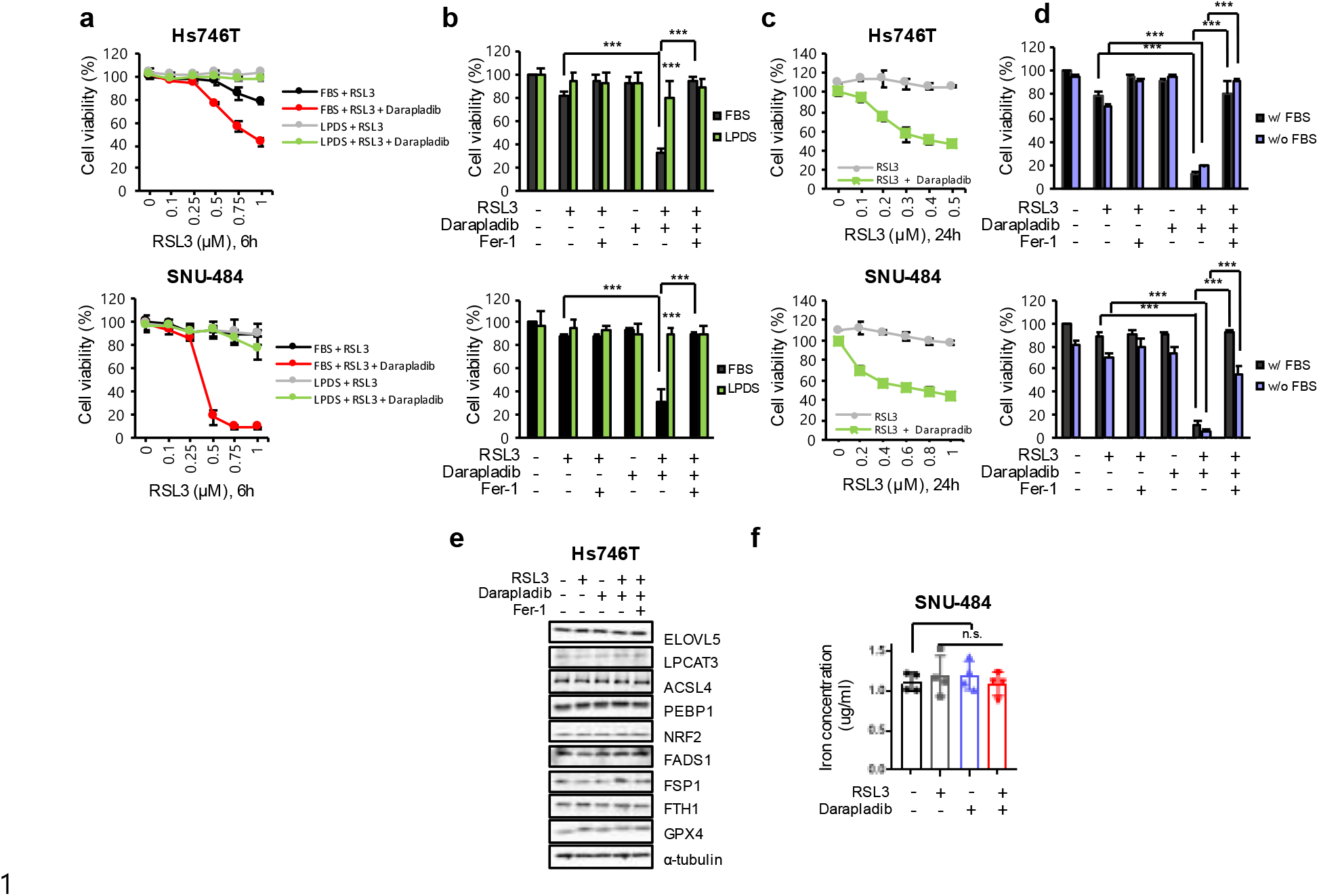
Darapladib is able to promote ferroptosis in the absence of lipoprotein. (a and b) Relative viability of Hs746T and SNU-484 cells treated with RSL3 and 2 μM darapladib cultured in medium containing FBS or LPDS for 6 h in the presence or absence of Fer-1. (c) Relative viability of cells in the indicated medium for 24 h. (d) Relative viability of cells treated with RSL3 and 2 μM darapladib in the presence or absence of FBS. (e) Western blots showing the expression levels of well-known ferroptosis regulators upon 0.2 μM RSL3 and 2 μM darapladib treatment as indicated. (f) Total iron level measured in the lysates of cells treated with 0.2 μM RSL3 and 2 μM darapladib. The data are the means ± SDs (n = 3 independent experiments).

### Darapladib does not affect known ferroptosis regulators or iron levels

We next explored the mechanism by which darapladib sensitizes cells to ferroptosis. First, we investigated the expression of well-known ferroptosis regulators, such as GPX4, NRF2, FSP1, ACSL4, LPCAT3, PEBP1, FADS1, and ELOVL5 (Lee *et al*., 2021b; Zhao et al., 2021). As previously reported, GPX4 bands were shifted due to covalent binding to RSL3 (Eaton et al., 2020), but the levels of GPX4 were unchanged (Fig. 3e). In addition, we observed no significant differences in protein expression upon RSL3 and/or darapladib treatment (Fig. 3e). Since the availability of free iron is also a key factor for ferroptosis execution (Chen et al., 2020), we also investigated free iron levels upon darapladib treatment. However, darapladib did not regulate intracellular iron levels (Fig. 3f).

Lp-PLA2 is primarily produced by monocytes/macrophages and catalyzes the hydrolysis of oxidized phospholipids (oxPLs), mainly PC on LDL, into lysophosphatidylcholine (LysoPC) and oxidized nonesterified fatty acids (oxNEFAs) (Karakas and Koenig, 2010). In particular, lysoPC induces ROS production, contributing to atherosclerosis and plaque destabilization (Chang et al., 2006; Portman and Alexander, 1969; Yamakawa et al., 2002). Therefore, Lp-PLA2 is regarded as a risk factor for cardiovascular disease, and darapladib (SB480848) was developed to specifically inhibit Lp-PLA2 to treat atherosclerosis, although darapladib recently failed in phase III clinical trials due to the lack of efficacy (O’Donoghue et al., 2014; White et al., 2014; Wilensky et al., 2008). As lysoPC has previously been shown to promote AA release from endothelial cells (Wong et al., 1998), we hypothesized that darapladib reduces the extracellular levels of lysoPC, thereby abolishing the export of AA and eventually facilitating ferroptosis. Unexpectedly, however, the levels of lysoPC and AA in the medium were slightly increased or remained unchanged upon darapladib treatment (Supplementary Fig. 5a). Interestingly, medium supplemented with FBS contained abundant lysoPCs, which decreased rapidly over time during incubation and were probably absorbed into cells and used for phospholipid synthesis through the Lands cycle (Kamphorst et al., 2013). While we cannot rule out the possibility that Lp-PLA2-mediated lyso-PCs production is still linked to ferroptosis, our findings point to the existence of a dominant mechanism by which Lp-PLA2 regulates ferroptosis independently of lysoPC production.

### Lp-PLA2 inhibits ferroptosis

Given that lipoproteins and extracellular lysoPCs are is dispensable for darapladib-induced ferroptosis, we wonder whether Lp-PLA2 is indeed associated with ferroptosis pathway. Therefore, we employed small interfering RNA (siRNA) pools containing four different siRNAs against Lp-PLA2, and efficient knockdown by the siRNA was confirmed at the mRNA level (Supplementary Fig. 6a). Consistent with darapladib, depletion of Lp-PLA2 also promoted RSL3-induced ferroptosis in both Hs746T and SNU-484 cells (Supplementary Fig. 6b).

Next, we established H1299 and YCC-16 cells deficient for *PLA2G7*, which encodes Lp-PLA2, using CRISPR-Cas9 systems and confirmed complete knockout at the mRNA level (Fig. 4a and Supplementary Fig. 7a). Again, *PLA2G7*-KO cells were more sensitive to RSL3 than parental cells, and the increased sensitivity to ferroptosis was reversed by Fer-1 (Fig. 4b-g and Supplementary Fig. 7b-f). In addition, the level of lipid peroxidation was increased in *PLA2G7*-KO cells (Fig. 4h and Supplementary Fig. 7g). Furthermore, under cysteine-deficient conditions, wild-type cell viability was not greatly reduced, but *PLA2G7*-KO cells underwent rapid cell death, which was attenuated by Fer-1 (Fig. 4i,j and Supplementary Fig. 7h). Similar to the findings with darapladib, *PLA2G7*-deleted cells showed no significant alterations in several key ferroptosis regulators (Supplementary Fig. 7i). Finally, *PLA2G7*-KO cells exhibited reduced sensitivity to darapladib-enhanced ferroptosis, supporting the requirement of Lp-PLA2 for darapladib (Fig. 4k and Supplementary Fig. 7j). These data imply that Lp-PLA2 produced in cells rather than in serum might play a key protective role in ferroptosis.

**Fig. 4.**
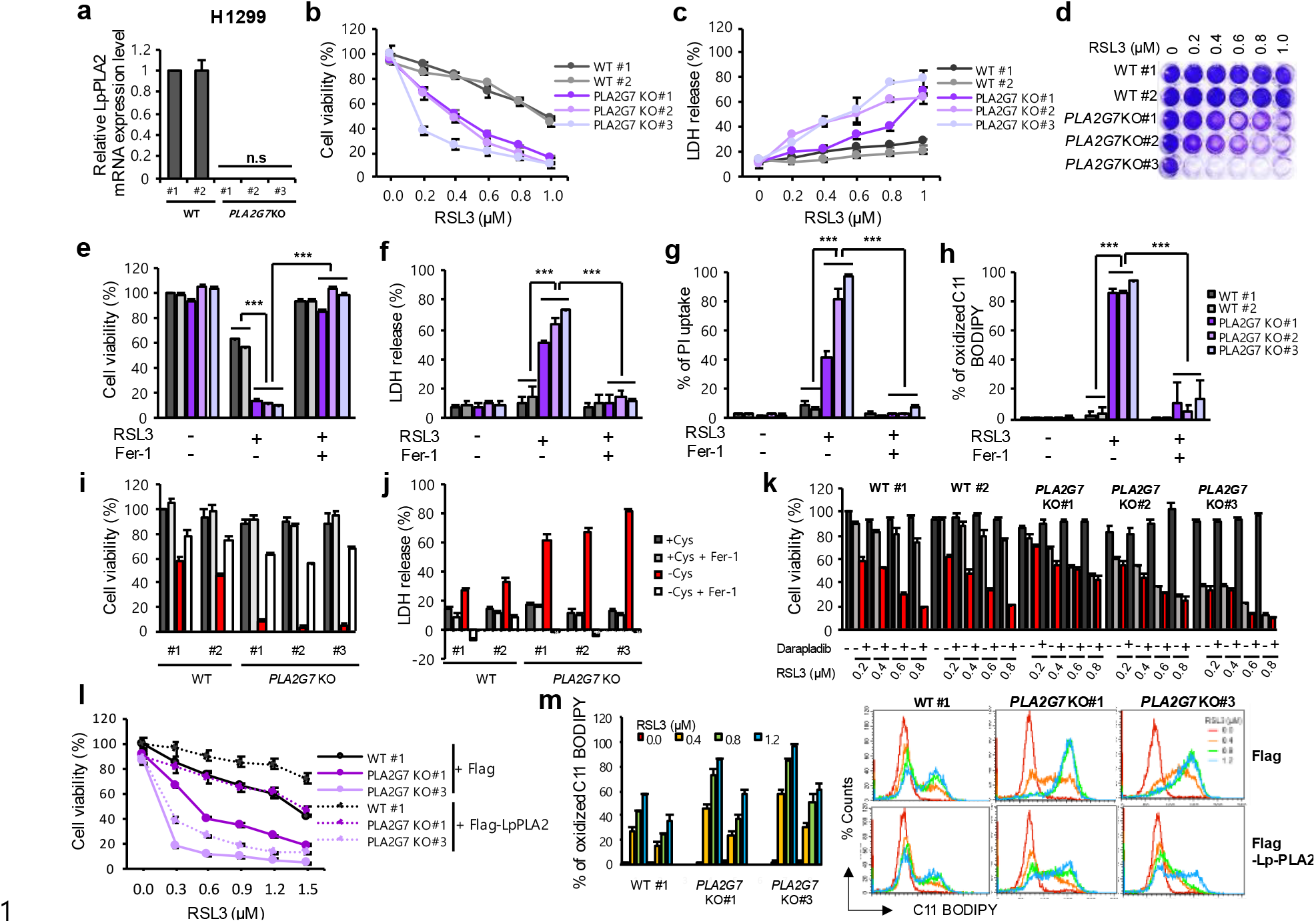
Lp-PLA2 negatively regulates lipid peroxidation and ferroptosis. (a) Analysis of mRNA expression in Lp-PLA2 depletion cells. Relative expression levels were normalized to the β-actin expression levels. (b and c) Relative cell viability and LDH levels in WT and *PLA2G7* KO H1299 cells treated with RSL3 for 20 h. (d) Crystal violet staining result after various concentration of RSL3 treatment in WT and *PLA2G7* KO H1299 cells. (e, f and g) Relative cell viability, LDH levels and PI uptake in WT and *PLA2G7* KO H1299 cells treated with RSL3 in the presence or absence of Fer-1 for 20 h. (h) Lipid peroxidation level in WT and *PLA2G7* KO H1299 cells treated with RSL3 for 1.5 h. (i and j) Cell viability and LDH level from WT and *PLA2G7* KO H1299 cells cultured with cysteine deficient medium for 48 h. (k) Relative cell viability from WT and *PLA2G7* KO H1299 cells treated with various concentration of RSL3 or/and 0.5 µM darapladib for 48 h. (l) Relative viability of WT and *PLA2G7* KO H1299 cells treated with RSL3 after ectopic expression of Lp-PLA2. (m) Lipid peroxidation level in WT and *PLA2G7* KO H1299 cells treated with RSL3 after ectopic expression of Lp-PLA2.

We next explored whether ectopic expression of Lp-PLA2 protein in cells could ameliorate RSL3-induced ferroptosis. Hs746T cells overexpressing Lp-PLA2 exhibited reduced susceptibility to RSL3-induced cell death (Fig. 4l and Supplementary Fig. 7k). In addition, overexpression of Lp-PLA2 reduced RSL3-induced lipid peroxidation (Fig. 4m and Supplementary Fig. 7l). Taken together, these findings suggest that Lp-PLA2 is a bona fide negative regulator of ferroptosis.

### Intracellular Lp-PLA2 is responsible for ferroptosis sensitization

While Lp-PLA2 is known to be secreted into the extracellular matrix and to interact with lipoprotein (Mallat et al., 2010), our data suggest that lipoprotein is not essential for darapladib sensitization of ferroptosis. In addition, several previous reports have implied that Lp-PLA2 is localized in the cytoplasm (Lehtinen et al., 2017; Vainio et al., 2011a; Xiao et al., 2012). We also observed a comparable amount of ectopic Lp-PLA2 in cell pellets compared to medium (Fig. 5a). In addition, overexpressed Lp-PLA2 existed in both the cytoplasm and plasma membrane (Fig. 5b).

**Fig. 5.**
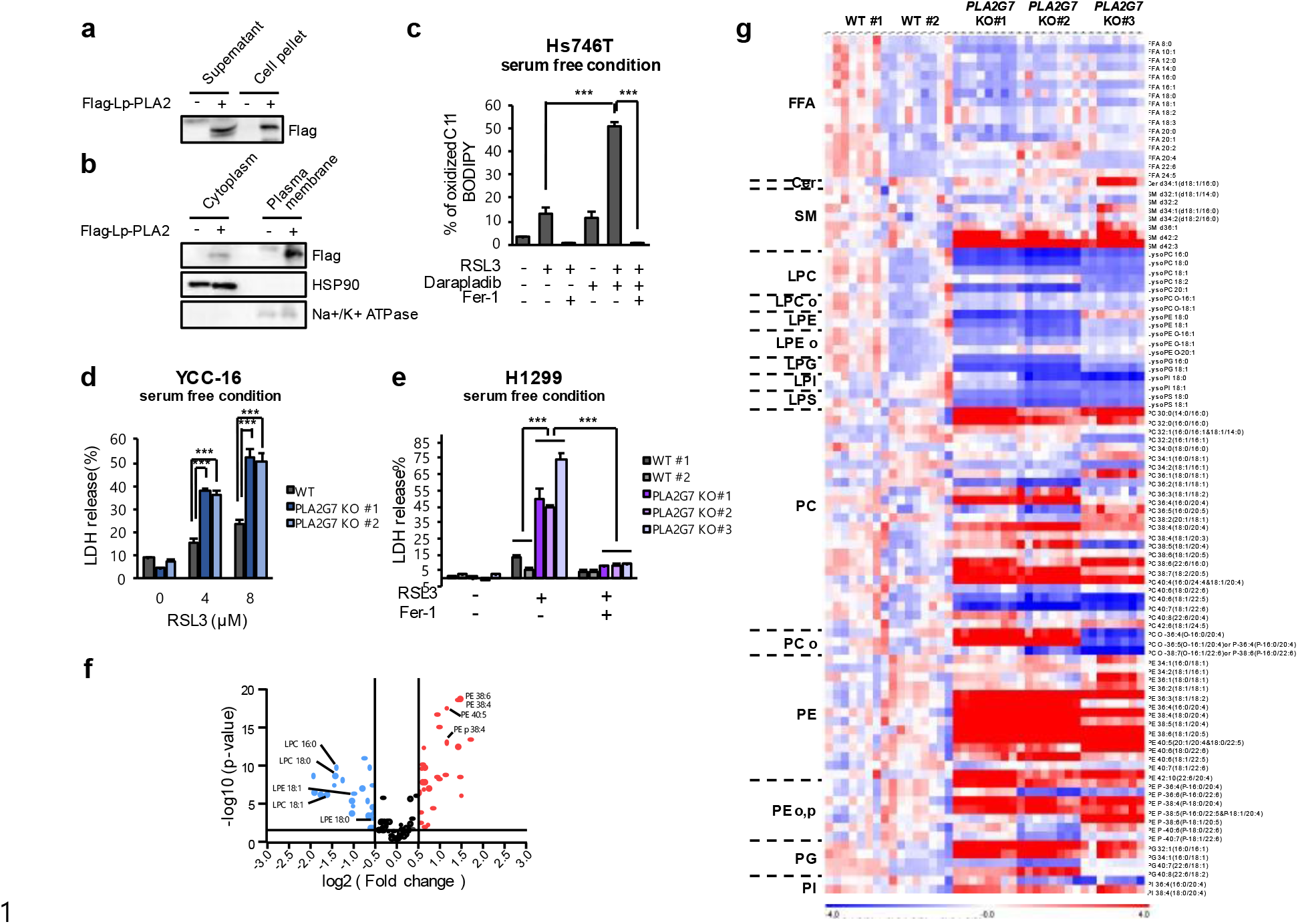
Darapladib might target intracellular Lp-PLA2, thereby promoting ferroptosis by inducing the accumulation of PE species. (a) Western blot analysis to confirm the presence of Lp-PLA2 in the pellet and supernatant after ectopic expression of Lp-PLA2 in Hs746T cells. (b) Western blot analysis to confirm the presence of Lp-PLA2 in the cytoplasm and plasma membrane after ectopic expression of Lp-PLA2 in Hs746T cells. (c) Lipid peroxidation levels in Hs746T cells treated with RSL3 or/and 2 μM darapladib for 0.5 h and cultured with FBS-deficient medium. Lipid oxidative potential was assessed by flow cytometry and microscopy using C11-BODIPY^581/591^. (d and e) LDH levels in WT and *PLA2G7* KO cells (YCC-16 and H1299 cells) treated with RSL3 and cultured with FBS-deficient medium for 24 h. (f) Volcano plot of lipid classes showing the log2(fold change) and –log10(p) values in control vs. *PLA2G7* KO H1299 cells. (g) Lipidomic analysis acquired from the UPLC/QTOF MS spectra of WT and *PLA2G7* KO H1299 cells. Each value in the heatmap is a coloured representation of a calculated Z score. Lipids are presented as the total number of carbon atoms and the total number of double bonds. Lipids with dissimilar MS/MS spectra but identical numbers of carbon atoms and an equal number of double bonds are marked with ^a^ and ^b^. n = 8 independent experiments. FFA: free fatty acid, Cer: ceramide, SM: sphingomyelin, lysoPC: lysophosphatidylcholine, lysoPE: lysophosphatidylethanolamine, lysoPG: lysophosphatidylglycerol, lysoPI: lysophosphatidylinositol, lysoPS: lysophosphatidylserine, PC: phosphatidylcholine, PE: phosphatidylethanolamine, PG: phosphatidylglycerol, PI: phosphatidylinositol, PS: phosphatidylserine.

To exclude the involvement of extracellular Lp-PLA2, the medium of the cells was replaced with serum-free medium to remove the existing Lp-PLA2 in the medium, and the cells were immediately stimulated with RSL3. Interestingly, darapladib was still able to increase lipid peroxidation levels at 30 min after RSL3 treatment (Fig. 5c). Similarly, Lp-PLA2-KO cells were more sensitive to ferroptosis than control cells in the presence of fresh medium without serum (Fig. 5d, e). These data suggest that intracellular Lp-PLA2 might have a protective role against ferroptosis.

### Deletion or inhibition of Lp-PLA2 induce lipid remodeling that is favorable to ferroptosis

Since intracellular PUFAs in phospholipids are crucial in ferroptosis sensitivity, we hypothesize that intracellular Lp-PLA2 might control lipid composition in cells. When lipidomic changes under *PLA2G7* deficiency were assessed using LC-MS, we found that the levels of PEs and PCs were generally increased in *PLA2G7* KO cells, while the levels of lysoPEs and lysoPCs were decreased in these cells (Fig. 5f,g and Supplementary Fig. 8a,b). The increase in PE and PC species with three or more double bonds was remarkable when compared to those with fewer double bonds (Fig. 5g). In particular, we also found that well-established pro-ferroptotic phospholipids such as PE-38:4 (18:0/20:4) are enriched in *PLA2G7* KO cells (Supplementary Fig. 8b).

We next tested whether the inhibition of Lp-PLA2 by darapladib results in the similar lipidomic changes observed in *PLA2G7* KO cells. Lipid profile analysis reveals that the levels of PEs were generally increased in darapladib-treated cells, while the levels of free fatty acids (FFAs) and lysoPEs were decreased by darapladib (Fig. 6a and Supplementary Fig. 9a-c). This lipidomic changes were observed as early as 1 h and lasted for 4 h, indicating that phospholipid remodeling, known as the Lands cycle, occurs very quickly within the cell. In addition to diacyl PE species, ether lipids such as PE plasmalogens (PE-p) species, which are crucial for ferroptosis, accumulated, while lysoPE-p species were reduced by darapladib (Cui et al., 2021; Zou et al., 2020) (Supplementary Fig. 9a-c). In addition, other phospholipids such as PI, PS, and PG, may be the target of Lp-PLA2 as these PLs and lysoPLs are oppositely regulated by Lp-PLA2 deficiency or inhibition (Fig 5 and Supplementary Fig. 9). Unlike *PLA2G7* KO cells, PC species, the most abundant phospholipids, remained unchanged in cells treated with darapladib (Supplementary Fig. 9a-c). These data suggest that darapladib induces the accumulation of AA and AdA containing PE or PE-p, rendering cells sensitive to ferroptosis.

**Fig. 6.**
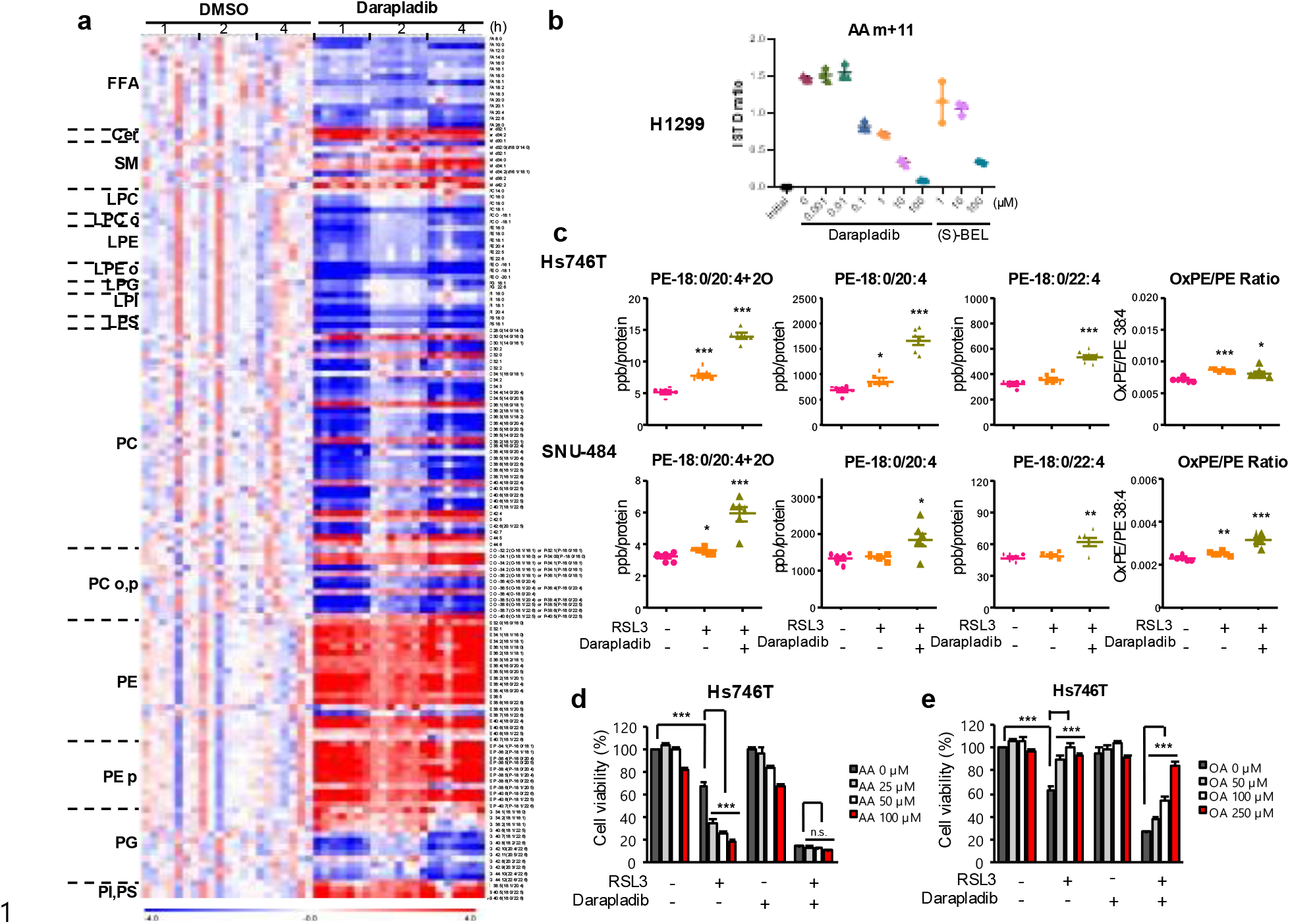
Darapladib induces global rearrangement in lipid compositions. (a) Lipidomic analysis acquired from the UPLC/QTOF MS spectra of Hs746T cells treated with 2 µM darapladib for 1, 2, and 4 h. Each value in the heatmap is a coloured representation of a calculated Z score. Lipids are presented as the total number of carbon atoms and the total number of double bonds. Lipids with dissimilar MS/MS spectra but identical numbers of carbon atoms and an equal number of double bonds are marked with ^a^ and ^b^. n = 7 independent experiments. (b) PE-18:0/20:4-d11 tracing to confirm Darapradib’s ability to block PE-18:0/20:4. (c) Concentrations of oxidized PE-C18:0/C20:4, PE-C18:0/C20:4, and PE-C18:0/C22:4 after treatment with RSL3 or/and 2 μM darapladib for 3 h in Hs746T cells and for 4 h in SNU-484 cells. The ratios of oxidized PE-C18:0/C20:4 to PE-C18:0/C20:4 are also shown. The levels of oxidized PE, PE-C18:0/C20:4, and PE-C18:0/22:4 were determined using LC–MS/MS. The concentrations were normalized to the cellular protein level. The data are the means ± SDs (n=6 independent experiments). (d and e) Relative viability of Hs746T cells pretreated with AA or OA for 20 h and treated with RSL3 or/and 2 μM darapladib for 20 h.

Since Lp-PLA2 is known to preferentially cleavage oxidized PC, we tested the ability of darapladib directly inhibit the cleavage of PE-18:0/20:4 by tracing PE-18:0/20:4-d11 and its cleavage product AA-d11. While cell lysates efficiently reduced PE-18:0/20:4 and produced AA-d11, darapladib strongly suppressed these responses (Fig. 6b). Furthermore, lysates from *PLA2G7* KO cells had a reduced ability to cleave PE-18:0/20:4-d11, indicating that Lp-PLA2 is responsible for PE cleavage (Fig. 6c).

Interestingly, the levels of saturated fatty acids and monounsaturated fatty acids (MUFAs), which are also generated by the de novo fatty acid synthesis pathway, were reduced by darapladib. Therefore, we evaluated the expression of several proteins, such as SREBP1, ACC1, and FASN. We confirmed that the precursor form of SREBP1 was reduced and that phosphorylation of ACC was significantly increased by darapladib treatment, suggesting that the de novo fatty acid synthesis pathway was generally inhibited (Supplementary Fig. 9d). Notably, the reduction in oleic acid (OA, C18:1), which can strongly suppress ferroptosis (Magtanong et al., 2019; Ubellacker et al., 2020), was much more pronounced than the reductions in saturated fatty acids, which may contribute to ferroptosis. These data imply that darapladib mediates the reduction in MUFA levels and accumulation of PE-PUFAs, thereby promoting ferroptosis.

Recently, Lp-PLA2 was shown to specifically cleave oxidized 1-stearoyl-2-arachidonyl-PE (SAPE; hereinafter referred to as PE-18:0/20:4), thereby abrogating engulfment of ferroptotic cells by macrophages (Tyurin et al., 2014). We therefore investigated whether darapladib also protects oxidized PE-18:0/20:4 (oxPE-18:0/20:4) from cleavage into PE and FFA by Lp-PLA2 upon ferroptosis induction. While the levels of oxPE-18:0/20:4 were slightly increased in response to a low concentration of RSL3 or darapladib, cotreatment with RSL3 and darapladib drastically increased the levels of oxPE-18:0/20:4 in both Hs746T and SNU-484 (Fig. 6c and Supplementary 10a). However, this induction was accompanied by increases in non-oxidized PE-18:0/20:4 and in PE-18:0/22:4, as supported by our targeted lipid analysis Nevertheless, the slight increase in the ratio of oxPE-18:0/20:4 to PE-18:0/20:4 by darapladib was also observed (Fig. 6c and Supplementary Fig. 10a), suggesting that Lp-PLA2 activity against oxidized SAPE also partially contributes to ferroptosis sensitivity. Consistently, *PLA2G7* KO cells exhibited a modest increase in oxPE-18:0/20:4 levels, possibly due to the increase in PE-18:0/20:4, RSL3 treatment significantly elevated oxPE-18:0/20:4 levels (Supplementary Fig. 10b).

We next used a docking model to predict binding affinity between Lp-PLA2 and various phospholipids in order to estimate the Lp-PLA2’s ability to access these phospholipids. Interestingly, our prediction model suggests that Lp-PLA2 has a higher affinity for PE and PC than for PLs with other heads. Furthermore, peroxidation on AA chain seems to partially inhibit the interaction of Lp-PLA2 and SAPE (Supplementary Fig. 10c). These data suggest that Lp-PLA2 inhibition renders cells sensitive to ferroptosis by increasing the levels of PE-18:0/20:4 and protecting oxPE-18:0/20:4 from cleavage.

We have previously shown that supplementation with AA ultimately increases PE-18:0/20:4 levels, thereby sensitizing cells to ferroptosis (Lee *et al*., 2020). As we suggest that darapladib induces ferroptosis by ultimately increasing the levels of PE-18:0/20:4, we wondered whether darapladib could no longer promote ferroptosis in the presence of a high concentration of AA. As expected, AA increased the susceptibility of cells to ferroptosis, but darapladib only slightly exacerbated ferroptosis in the presence of AA (Fig. 6d). In contrast, supplementation with OA, the levels of which were decreased upon darapladib treatment, suppressed ferroptosis in the presence of RSL3 and darapladib (Fig. 6e). These data support the idea that both the increase in PE-18:0/20:4 and the decrease in OA are responsible for ferroptosis sensitization by darapladib.

### PLA2G7 is upregulated in various types of tumors and its inhibition with PACMA31 effectively suppresses gastric tumor growth

Although Lp-PLA2 encoded by PLA2G7 genes is known to be mainly expressed in monocytes and macrophages (Rosenson and Stafforini, 2012), its overexpression in tumors including prostate, colon, and breast cancer, has been reported in a number of studies (Huang et al., 2020; Lehtinen *et al*., 2017; Tannock et al., 2018; Vainio et al., 2011b; Wang et al., 2021b; Xu et al., 2013). We therefore investigated the expression of Lp-PLA2 in tumors compared to normal tissues in The Cancer Genome Atlas (TCGA) and found that Lp-PLA2 is upregulated in several types of cancer, including gastric cancer, esophageal cancer, lung adenocarcinoma, and breast cancer (Supplementary Fig. 11a). Since we focused on the function of Lp-PLA2 in gastric cancer, we further analysed the expression of Lp-PLA2 in several independent gastric cancer cohorts and observed the general overexpression of Lp-PLA2 in tumors compared to normal tissues (Supplementary Fig. 11b).

We next wondered whether darapladib also contributes to the anti-tumor effect upon ferroptosis induction.We first established H1299 cell lines stably expressing lentiviral shRNA for GPX4, which was selected in the presence of Fer-1 to prevent ferroptosis upon GPX4 depletion (Supplementary Fig. 12a). Removal of ferrostain-1 can induce lipid peroxidation and ferroptotic cell death, which were reinforced by darapladib, confirming that darapladib sensitized GPX4 depletion-induced ferroptosis (Fig. 7a,b and Supplementary Fig. 12b,c).

**Fig. 7.**
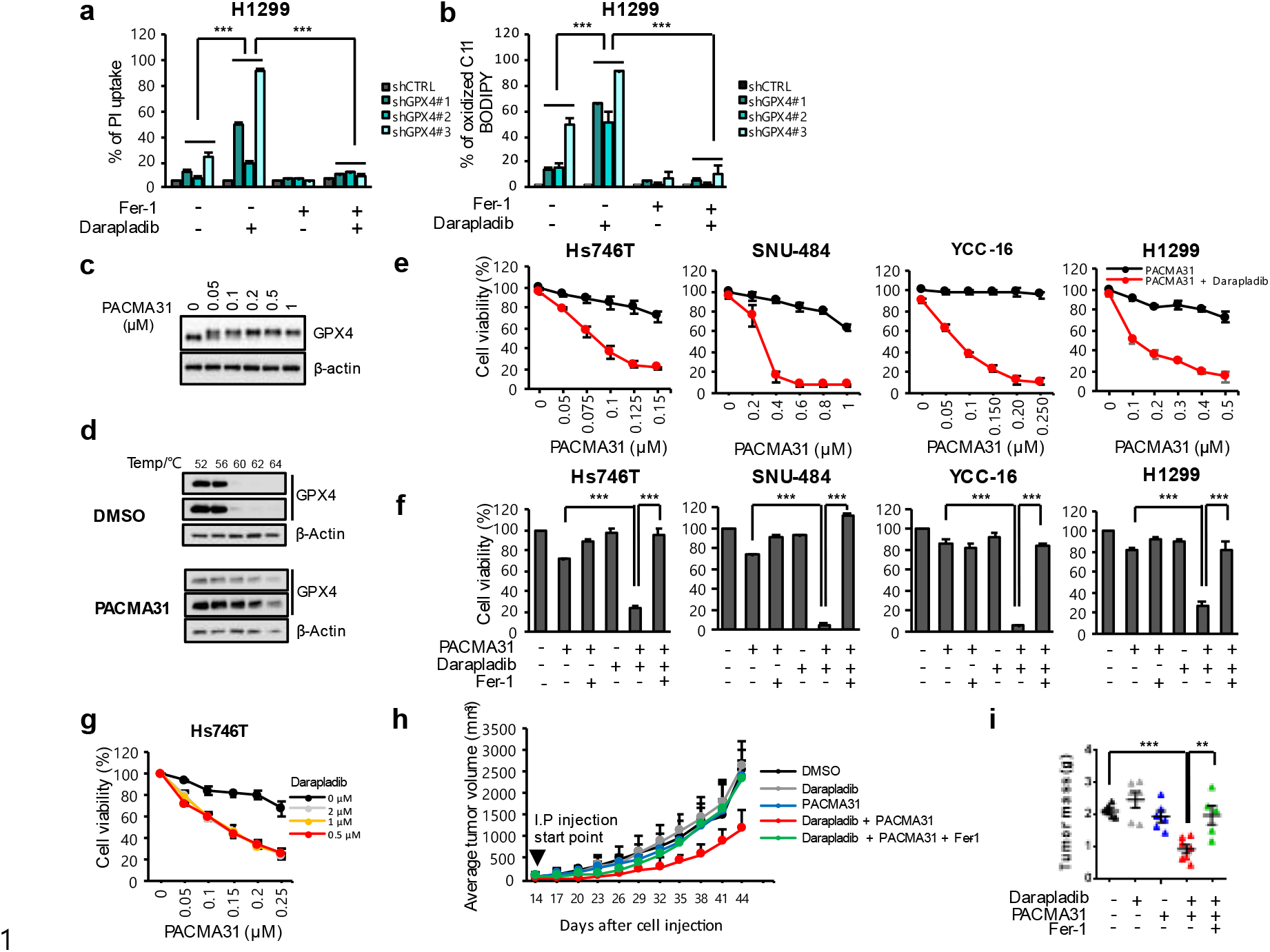
Darapladib enhances the anti-tumor activity of PACMA31 by accelerating ferroptosis. (a) PI uptake from WT and GPX4 depleted H1299cells treated with 5 uM darapladib in the presence or absence of 2 uM Fer-1 for 20 h. (b) Lipid peroxidation level in WT and GPX4 depleted H1299 cells treated with 5 uM darapladib in the presence or absence of 2 uM fer-1 for 1 h. (c) Western blot showing the band shift of GPX4 after PACMA31 treatment. (d) Thermal shift assay showing the stability of GPX4 in lysate of cells treated with PACMA31. (e and f) Relative viability of Hs746T, SNU-484, YCC-16 and H1299 cells treated with PACMA31 and/or 2 µM darapladib in the presence or absence of Fer-1 for 20 h. (g) Relative viability of Hs746T cells treated with PACMA31 in the presence of increasing concentrations of darapladib. The data are the means ± SDs (n = 3 independent experiments). (h and i) SNU484 cells were injected subcutaneously into immunodeficient nude mice to induce tumor formation. Eleven days after inoculation, the mice were treated with PACMA31 and/or darapladib. Intraperitoneal injections were carried out every 3 days as indicated. Each treatment group consisted of 5-7 mice treated with vehicle, 10 mg/kg PACMA31, 10 mg/kg darapladib, 10 mg/kg PACMA31 + 10 mg/kg darapladib or 10 mg/kg PACMA31 + 10 mg/kg darapladib + 10 mg/kg fer-1 for 28 days. The size of the tumor was measured every 3 days, and after approximately 4 weeks, the tumor was excised to measure the actual size and weight of the tumor.

Since most GPX4 inhibitors, including RSL3 and ML210, are not suitable for in vivo use (Viswanathan *et al*., 2017), we employed PACMA31, a previously known PDI inhibitor that was recently identified as a new GPX4 inhibitor and is available for in vivo study (Xu et al., 2012; Yan et al., 2021). We first confirmed that PACMA31 induced a band shift of GPX4 in a dose-dependent manner similar to other GPX4 inhibitors and increased thermal stability of GPX4 in cell lysates, suggesting the direct binding between PACMA31 and GPX4 (Fig. 7c, d). In addition, PACMA31 exhibited IC50 values comparable to those of other GPX4 inhibitors in cells, which was completely rescued by ferrostain-1, supporting PACMA31 is a reliable ferroptosis inducers (Supplementary Fig. 13a, b). Strikingly, PACMA31 killed cells synergistically with darapladib (Fig. 7e-g).

We then tested combination treatment of PACMA31 and darapladib in a mouse xenograft model established using SNU-484 cells. SNU-484 cells were injected subcutaneously into nude mice, after which the mice were treated with PACMA31 and/or darapladib. Treatment of mice with PACMA31 or darapladib alone showed no statistically significant difference compared to treatment with the vehicle control (Fig. 7h). However, tumor growth in mice treated with both PACMA31 and darapladib was drastically reduced compared to that in the other groups. Eventually, the size and weight of the tumors were significantly reduced in the combination treatment group compared to the other groups (Fig. 7i). Furthemore, ferrostain-1 rescued the reduced tumor growth and weight upon darapladib and PACMA31 treatment, suggesting that ferroptosis contribute to the tumor suppression (Fig. 7h, i). When we investigated lipid profiles in xenografted tumor treated with PACMA31 and/or darapladib, we observed a general increase in PE species in darapladib-treated xenograft tumor tissues, but not as pronounced as in the cells treated with darapladib (Supplementary Fig. 13c). Taken together, these findings suggest that inhibition of both Lp-PLA2 and GPX4 synergistically inhibits tumor growth in vivo.

## Discussion

The critical roles of ferroptosis in various human diseases including neurological disorders, ischemia-reperfusion injury, kidney damage, and blood disorders, have become widely recognized through extensive in vitro and in vivo studies (Jiang et al., 2021b; Ko et al., 2021; Lane et al., 2021; Wu et al., 2021; Zhang et al., 2021a; Zou *et al*., 2020). In addition, cancer cells, especially chemoresistant or persister cancer cells, are highly vulnerable to ferroptosis inducers, which suggests a strategy for anti-cancer therapy using ferroptosis (Hangauer *et al*., 2017; Lee *et al*., 2020; Rodriguez *et al*., 2022; Viswanathan *et al*., 2017). Most cells implement several protection mechanisms against ferroptosis: for example, the GPX4/GSH system primarily removes lipid peroxides, acting as a first line of defence. Therefore, inhibition of the GPX4/GSH system is necessary to induce ferroptosis in most contexts. NRF2 pathways generally suppress ferroptosis in various contexts (Dodson et al., 2019; Sun et al., 2016). In addition, FSP1/AIFM2 and DHODH regenerate CoQ10, allowing CoQ10 to directly neutralize lipid peroxy radicals (Bersuker et al., 2019; Doll et al., 2019; Mao et al., 2021). Furthermore, iPLA2β can directly cleave oxidized SAPE (HpETE) (Beharier *et al*., 2020; Chen *et al*., 2021; Sun *et al*., 2021). Finally, the endosomal sorting complexes required for transport (ESCRT) machinery ultimately protects against cell membrane destruction upon ferroptosis (Dai et al., 2020; Pedrera et al., 2021).

While AA and AdA are the most susceptible PUFAs to peroxidation, how these lipids are controlled in cells is largely unknown (Aldrovandi *et al*., 2021). The most abundant PUFA in serum and plasma is LA (C18:2), and a comparable amount of AA is also found in serum (Else, 2020). However, very long-chain PUFAs (VLC-PUFAs) with more than 22 carbons, such as AdA (C22:4), are rarely present in serum and plasma. In this regard, we recently reported that cell-autonomous synthesis of AA and AdA by fatty acid desaturases (FADSs) and elongases (ELOVLs) is critical for ferroptosis, although AA can be imported when PUFA synthesis is inhibited (Lee *et al*., 2020). Nevertheless, a number of studies have suggested that AA import and AA esterification by ACSL4 play a pivotal role in ferroptosis (Doll et al., 2017b; Liao et al., 2022; Tousignant et al., 2020). Therefore, precise studies are needed to determine how much each pathway contributes to the intracellular levels of AA under various conditions. In this study, we suggest that cell-autonomous recycling of AA by Lp-PLA2 is a crucial pathway that determines the amount of AA-containing PE and PE-p, thereby contributing ferroptosis resistance. In addition, this pathway might be favorable for a rapid response to ferroptotic stimuli, as PUFA synthesis or uptake seems to be a relatively slow process compared to PUFA recycling by Lp-PLA2.

While PLA2 superfamily members have the same fundamental function, their functions may differ in different contexts, such as in different subcellular locations and tissues. Lp-PLA2 is unique in that it mainly binds to lipoproteins outside cells and acts on phospholipids in lipoproteins. The most well-defined role of Lp-PLA2 is its contribution to atherosclerosis. Lp-PLA2 is strongly expressed in the necrotic core and cleaves oxLDLs, mostly oxPC, into lysoPCs and NEFAs, leading to macrophage death and thereby contributing to atherosclerosis (Kolodgie et al., 2006). However, apart from its classical role, Lp-PLA2 seems to suppress ferroptosis independently of lipoprotein, as Lp-PLA2-KO cells or cells treated with the Lp-PLA2 inhibitor darapladib were more sensitive to ferroptosis under lipoprotein-deficient or serum-deficient conditions (Fig. 3 and 5). In addition, the Lp-PLA2 inhibitor did not reduce the extracellular levels of lysoPC (Supplementary Fig. 5). Although we could not completely exclude the possible off-target effect of darapladib on the promotion of ferroptosis, the observation that Lp-PLA2 depletion or deletion also augments ferroptosis supports the idea that Lp-PLA2 is an important regulator of ferroptosis.

Lp-PLA2 not only cleaves oxidized LDL but also cleaves oxidized PS, an eat-me signal of apoptotic cells, thereby restricting phagocytosis by macrophages (Tyurin *et al*., 2014). A recent study has suggested that Lp-PLA2 also recognizes and cleaves oxPE-(C18:0/C20:4) upon ferroptosis stimuli, implying that oxPE-(C18:0/C20:4) acts as an eat-me signal of ferroptotic cells that is recognized by macrophages (Luo et al., 2021). However, whether these activities are related to ferroptotic cell death has not been studied. We also observed that the ratio between oxPE-(C18:0/C20:4) and PE-(C18:0/C20:4) was slightly increased by the inhibition of Lp-PLA2 (Fig. 6 and Supplementary Fig. 10). However, we also observed global rearrangement in the lipid profile upon the inhibition of Lp-PLA2. In particular, Lp-PLA2 seemed to exert PLA2 activity, as several phospholipids, such as PE, PI, and PS, accumulated while their corresponding lysophospholipids were downregulated upon Lp-PLA2 inhibitor treatment. Interestingly, PC, the most abundant phospholipid, was not significantly affected by the Lp-PLA2 inhibitor. These observations raise the possibility that Lp-PLA2 may also affect the Kennedy pathway, which is involved in the conversion of PC from PE. However, we observed these lipidomic changes within 1 hour, which seemed to be insufficient time for the lipidome to be changed through the Kennedy pathway. Rather, lysoPCs abundant in the medium were rapidly scavenged by the cells and used for PC synthesis, thereby contributing to maintaining intracellular PC levels under normal conditions. Although Lp-PLA2 inhibition may protect from PC cleavage in cells, it may also delayed PC synthesis, as the reduction of lysoPCs in the medium was slightly inhibited by darapladib (Supplementary Fig. 5). In addition to PE species, OA, the most abundant MUFA in serum that prevents ferroptosis, was downregulated by darapladib treatment, possibly contributing to the increased susceptibility to ferroptosis. Interestingly, while we did not observe drastic changes in lysoPC in the medium, supplementation with lysoPCs, such as lysoPC16:0 and lysoPC18:1, abrogated RSL3-induced ferroptosis, although the underlying mechanism requires further investigation (Supplementary Fig. 14).

As PLA2 is known to specifically cleave phospholipids at the sn-2 position into FFAs and lysophospholipids (lysoPLs) (Dennis et al., 2011), other PLA2 isoforms may control ferroptosis by regulating PUFA-containing PL abundance. While depletion of several PLA2 isoforms showed context-dependent promotion of ferroptosis, inhibition of each PLA2 isoforms with their inhibitors has no obvious effect on ferroptosis in cancer cells (Supplementary Fig. 15a, b). Interestingly, (S)-BEL, an inhibitor of iPLA2, is itself toxic to H9c2 cells at high concentration, but it slightly enhances ferroptosis (Supplementary Fig. 15c). Notably, darapladib sensitized various cell lines including H9c2 cells to ferroptosis, suggesting that Lp-PLA2 may be a common regulator of ferroptosis that conserving PEs containing PUFAs (Supplementary Fig. 15).

Induction of ferroptosis is an attractive strategy for cancer therapy, as a single inhibitor of GPX4 or SLC7A11 can drastically induce ferroptosis in cells. However, with its poor pharmacokinetic profile, the use of RSL3 in vivo is very limited, and only the anti-tumor activity of RSL3 in vivo has been tested by intratumoral injection as a proof-of-concept (Yang et al., 2014). In this study, we employed PACMA31, a recently identified GPX4 inhibitor that was previously known as a PDI inhibitor (Xu *et al*., 2012; Yan *et al*., 2021). We also confirmed that PACMA31 has high in vitro potency to kill cancer cells and directly binds to GPX4, as evidenced by a band shift. In combination treatment, PACMA31 and darapladib exhibited strong anti-tumor activity in a mouse xenograft model. Since GPX4 inhibition may be toxic to various normal tissues, including the heart, kidneys, and liver, how to deliver drugs specifically to cancer tissues remains a challenge. Since there were no toxicity issues in phase III clinical trials, darapladib may be used as a repositioning drug for cancer therapy. In summary, Lp-PLA2 reduces the levels of PUFA-containing PEs and cleaves oxidized PEs, thereby suppressing ferroptosis. Therefore, inhibition of Lp-PLA2 using darapladib may be beneficial for ferroptosis-inducing strategies for cancer therapy. Our findings provide new insights into possible combination strategies targeting both Lp-PLA2 and ferroptosis in cancer.

## Materials and Methods

### Cell culture

Hs746T, SNU-484, and YCC-16 cells were purchased from the KCLB (Korean Cell Line Bank) at Seoul National University and H1299, A549, HepG2 and H9c2 cells were purchased from the American Type Culture Collection (USA). The Hs746T, H1299, A549, HepG2 and H9c2 cells were cultured in Dulbecco’s modified Eagle’s medium (DMEM) (HyClone). The SNU-484 and H1299 cells were cultured in RPMI medium (Corning). The YCC-16 cells were cultured in minimal essential medium (MEM) (HyClone). All media were supplemented with 10% fetal bovine serum (Gibco) and 100X antibiotics (Gibco). All of the cells were incubated at 37 °C in a humidified atmosphere with 5% CO_2_.

### Western blotting and antibodies

Cells were lysed using Nonidet-P40 lysis buffer (1% NP-40, 150 mM NaCl, 10% glycerol, 1 mM EDTA, and 20 mM Tris-HCl [pH 7.4]). Then, the membranes were blocked with 5% skim milk in TBS-Tween 20 (TBST), incubated with primary antibodies overnight at 4 °C and then incubated with secondary antibodies for 1 h at room temperature. For detection of ELOVL5 protein, samples were not boiled as previously described (Lee *et al*., 2020). The primary antibody against ACC (3662), ACC-p (3661), FASN (3180), S6K (9202) and S6K-p (9205) were purchased from Cell Signaling Technology (USA). SREBP1 (557036) was purchased from BD Biosciences (USA). ELOVL5 (sc-398653), ACSL4 (sc-365230), FSP1(sc-377120), PEBP1 (sc-376925) and HSP90 (sc-13119) were purchased from Santa cruz (USA). GPX4 (ab125066) and Na/K ATPase (ab254025) were purchased from Abcam (UK). NRF2 (16396-1-AP) and FADS1 (10627-1-AP) were purchased from Proteintech (USA). LPCAT3 (16-999) was purchased from prosci (USA). Flag (F3165), Actin (A5316) and a-tubulin (T9026) were purchased from Sigma-Aldrich (USA).

### Reagents

Darapladib (S7520), Fer-1 (S7243), RSL-3 (S8155), Erastin (S7142) and zVAD-fmk (zVAD, S7023) were purchased from Selleck Chemicals (USA). PACMA31 (5116) was purchased from Tocris (UK). Liproxstatin-1 (SML1414), LDL (L7914), HDL (L8039), VLDL (437647), ML210 (SML0521), Lipoprotein Deficient Serum (S5519) were purchased from Sigma–Aldrich (USA). Nec-1 (BML-AP309) was purchased from Enzo Life Sciences. MAFP (70660), (S)-BEL (70700), MJ33 (90001844), JKE1674 (30784), AA (90010) and OA (90260) were purchased from Cayman Chemical (USA).

pCMV3-SP-N-FLAG-Lp-PLA2 (HG10848-NF) was purchased from Sinobiological (China) and was subcloned into pCMV2-FLAG vector.

### Cell viability and LDH release assays

Cell viability was determined by measuring the cellular ATP levels using CellTiter-Glo reagent (CellTiter-Glo® 2.0 Assay, Promega) according to the manufacturer’s protocol. Cells were seeded at a density of 3 × 10^4^ cells/well in 200 µl of culture medium in 48-well plates. For a low density, 2 × 10^4^ cells/48 well were seeded in 48-well plates. The day after seeding, chemicals were administered at the concentrations and times indicated in the text. A volume of solution equal to that of the culture medium was added, and the cells were incubated at room temperature for 30 minutes. Then, 150 µl of the supernatant was transferred to a 96-well plate, and the absorbance was measured at 450 nm using a microplate reader. Cell-free supernatants were collected and assessed via LDH release assay with a Cytotoxicity Detection Kit (Roche). The percent LDH release was calculated according to the manufacturer’s instructions.

### Crystal violet staining

A total of 1 × 10^5^ cells were cultured in 24-well plates and exposed to drugs for 20 h. After the medium was removed, the cells were washed in cold phosphate-buffered saline (PBS), fixed with 100% methanol, and stained with 0.25% crystal violet solution.

### Cysteine deprivation

For cysteine deprivation, cysteine-free DMEM (lacking glutamine, methionine, and cysteine; Gibco) supplemented with methionine, 10% dialyzed FBS (26400044; Gibco) and 2 mM L-glutamine (Gibco) was used. Cells were washed with PBS and cultured with fresh DMEM or cysteine-free DMEM for 18 h in the presence or absence of Fer-1.

### Lipid peroxidation assay

Lipid peroxidation was measured with C11-BODIPY (581/591) (BODIPY™ 581/591 C11, Invitrogen). When lipid peroxidation increases, the fluorescence shifts from red to green. Cells were treated with RSL3 and/or darapladib and incubated for the indicated times. Next, 2.5 µM C11-BODIPY (581/591) was added, and the cells were incubated at 37 °C for 15 min. The cells were harvested by washing twice with PBS, transferred to a 1.5 ml microfuge tube (3000 rpm, 3 min), and then resuspended in 0.5 ml of PBS for flow cytometry. The cells were analyzed using the 488-nm laser of a flow cytometer (BD FACS Calibur) for excitation, and data were collected from the FL1 detector. A minimum of 10,000 cells were analyzed per condition.

### Iron assay

The iron concentration was determined using an iron colorimetric assay kit (Sigma). According to the manufacturer’s instructions, cells were harvested after washing with ice-cold PBS and homogenized in 5X volumes of Iron Assay Buffer and centrifuged at 16,000 × g for 10 min at 4°C to obtain the supernatant. 5 μL of Iron Reducer was added to each of the sample (50 µl/well) wells to reduce Fe3+ to Fe2+ and incubated for 30 min at 25°C. Subsequently, 100 μL of Iron Probe to each well and incubated the reaction in the dark for 60 min at 25°C. The absorbance was detected at 593 nm using microplate reader. Absorbance values were calibrated to a standard concentration curve to calculate the concentration of iron.

### RNAi-mediated gene knockdown

An ON-TARGET plus SMARTpool for human PLA2G7 siRNA (L-004903-00-0020), PA2G4A siRNA (L-0009886-00-0005), PLA2G2A siRNA (L-009901-00-0005), PLA2G6 siRNA (L-009085-00-0005) and a nontargeting pool (siNT, D-001810-10) were purchased from Dharmacon (USA). And An ON-TARGET plus individual set for human PLA2G7 siRNA (LQ-004903-00-0005) were purchased from Dharmacon (USA). Cells were transfected with 20 nM siRNA using Lipofectamine RNAiMAX (Invitrogen) according to the manufacturer’s protocol.

### Lp-PLA2 KO in H1299 and YCC-16 cell lines

The LP-PLA2-KO cell lines were produced by homologous recombination with the CRISPR–Cas9 system. The CRISPR–Cas9 constructs of PLA2G7 (LP-PLA2) were made by ligation of double-stranded antisense/sense oligos to the short guide RNAs (sgRNAs) of interest in pSpCas9(BB)-2A-GFP (PX458; Addgene #48138) and pSpCas9(BB)-2A-RFP plasmids. The CRISPR vector was created based on a manufacturer’s protocol. The CRISPR plasmid was cotransfected into cells using Lipofectamine 3000 transfection reagent (L3000015, Invitrogen). Forty-eight hours following transfection, green and red fluorescent protein-positive cells were selected by FACS and seeded into 96-well plates, and single clones were expanded. The insert sequences of the constructs are as follows:

sgRNA1-PLA2G7-S: CACCGGACCAATCTGCTGCAGAAAT, sgRNA1-PLA2G7-AS: AAACATTTCTGCAGCAGATTGGTCC, sgRNA2-PLA2G7-S: CACCGGCAGTAATTGGACATTCTTT, and sgRNA2-PLA2G7-AS: AAACAAAGAATGTCCAATTACTGCC.

### Total RNA extraction and RT–qPCR

Total RNA was extracted using TRIzol (Gibco) according to the manufacturer’s protocol. First-strand complementary DNA (cDNA) synthesis was performed using 2 μg of total RNA with M-MLV reverse transcriptase (Promega) according to the manufacturer’s protocol. All RT–qPCR was performed with SYBR Premix Ex Taq^TM^ (Takara Bio Inc., Japan) using a CFX96^TM^ Real-Time System (Bio–Rad Laboratories, USA). The primer sequences were as follows: Lp-PLA2 (forward), 5′-TGGCTCTACCTTAGAACCCTGA -3′; Lp-PLA2 (reverse), 5′-TTTTGCTCTTTGCCGTACCT-3′; β-Actin (forward), 5′-CTGGCACCCAGCACAATG-3′; and β-Actin (reverse), 5′-GCCGATCCACACGGAGTACT-3′.

### Thermal shift assay

Thermal shift assay was carried out in the lysate of cells treated with DMSO or PACMA31 using a BioRad real time PCR instrument. Temperature was increased from 52 to 62 °C. After that, western blot was performed to confirm the stability of GPX4.

### Lipidomic profiling using UPLC/QTOF MS

For global lipid profiling, ultra-performance liquid chromatography (UPLC) coupled with quadrupole time-of-flight mass spectrometry (QTOF-MS) was used to analyze the lipids in cells and culture media. For intracellular lipid analysis, 3×10^5^ cells were washed twice with PBS and extracted by scraping into 300 μl of 40:40:20 (v/v) methanol:acetonitrile:water containing 0.5% formic acid. The lipid extracts were cleared by centrifugation at 12,700 rpm for 15 min at 4 °C, and the supernatants were transferred to vials for lipidomic analysis. To measure the lipids in the culture media, the supernatant of each well was transferred to a 1.5 ml EP tube. Extracellular lipids were extracted by adding 800 μl of 50:50 (v/v) methanol:acetonitrile containing 0.625% formic acid to 400 μl of culture medium and centrifuging the mixture. Eight hundred microlitres of supernatant was transferred and dried under nitrogen gas. The medium extract was reconstituted in 80 μl of a 40:40:20 (v/v) methanol:acetonitrile:water solution. Tridecanoic acid and SPLASH Lipidomix (Avanti Polar Lipids, USA) were used as internal standards. Lipid analysis was conducted using an ACQUITY UPLC system (Waters, Milford, MA, USA) coupled with a triple TOF1™ 5600 mass spectrometer equipped with an electrospray ionization (ESI) source (AB Sciex, Concord, ON, Canada). Chromatographic separation of lipids in the cells and media was performed using an ACQUITY UPLC CSH C18 column (2.1 mm × 100 mm, 1.7 μm; Waters) at 55 °C and a flow rate of 0.4 mL/min. The binary mobile phases comprised 10 mM ammonium acetate in water:acetonitrile (60:40 v/v, solvent A) and isopropanol:acetonitrile (90:10 v/v, solvent B). The gradient was as follows: 40-43% B from 0-2 min, 43-50% B from 2-2.1 min, 50-54% B from 2.1-12 min, 54-70% B from 12-12.1 min, 70-99% B from 12.1-18 min, 99-40% B from 18-18.1 min, and 40% B from 18.1-20 min to equilibrate. The mass spectrometer was operated in negative ion mode, and the mass range was set at 80-1500 m/z for analysis of the lipid extracts. Total ion chromatograms were acquired using the following operating parameters: a capillary voltage of -4,500 V, a nebulizer pressure of 50 psi, a drying gas pressure of 60 psi, a curtain gas pressure of 30 psi, a source temperature of 500 °C, a declustering potential of -90 eV, a collision energy of -10 eV for single MS, and a collision energy of -45 eV for MS/MS. Data from MS/MS analyses were acquired using automatic fragmentation, in which the five most intense mass peaks were fragmented. Mass accuracy was maintained with an automated calibrant delivery system interfaced to the second inlet of the DuoSpray ion source. All the samples were pooled in equal volumes to generate quality control (QC) samples, which were analyzed prior to sample acquisition and after every 7 samples. Spectral data were preprocessed using MarkerView^TM^ software (AB Sciex) and normalized to the total spectral area. Lipid metabolites were identified using online databases (DBs; HMDB, METLIN, and LIPID MAPS). Identification was confirmed using the MS/MS patterns and retention times of lipid standard compounds.

### Analysis of AA and lysoPCs in medium

Extracellular lipid extracts were analyzed by UPLC/triple-quadrupole mass spectrometry (UPLC/TQ MS) in multiple reaction monitoring (MRM) mode using mobile phases and a column in the same manner as that for lipidomic profiling. An Agilent 1290 Infinity II LC and Agilent 6495 Triple Quadrupole MS system equipped with an Agilent Jet Stream ESI source (Agilent Technologies) was used for the analysis. Mass Hunter Workstation (ver. B.06.00, Agilent Technologies) software was used for data acquisition and analysis. Samples were eluted at 0.2 mL/min for 22 min. Gradient elution was conducted as follows: 40-55% B from 0-5 min, 55-60% B from 5-11 min, 60-99% B from 11-14 min, 99% B from 14-18 min, 99-40% B from 18-18.1 min, 40% B from 18.1-22 min. MS/MS experiments were conducted in negative ion mode. The MS system was operated using the following parameter settings: a gas temperature of 220 °C, a nebulizer gas of nitrogen at 30 psi, a sheath gas temperature of 300 °C, and a sheath gas flow rate of 11 L/min.

### Oxidized PE and PE analysis in cells

For oxidized PE, PE-(18:0/20:4), and PE-(18:0/22:4) analysis in cells, the lipids were extracted in 500 μl of 80:20 (v/v) methanol:water and 800 μl of chloroform. The lipid extracts were dried under nitrogen gas and reconstituted in 50:25:25 (v/v/v) isopropanol/acetonitrile/water containing PE-(33:1-d7) (Sigma–Aldrich, USA) as an internal standard. Targeted analysis of lipids was performed on an ACQUITY UPLC I-Class PLUS (Waters, Milford, MA, USA) and a Xevo TQ-XS Triple Quadrupole MS system (Waters, Milford, MA, USA) in MRM mode. MassLynx (Waters, Milford, MA, USA) software was used for data acquisition and analysis. The mobile phases and chromatographic separation were the same as those used for AA and lysoPC analysis. Individual standards were used for optimization of the conditions for each metabolite and for preparation of a series of calibration solutions to generate calibration curves.

### TCGA Pan-Cancer data and microarray data for gastric cancer analysis

Publicly available data from The Cancer Genome Atlas (TCGA) data were analyzed in the current study. Clinical information and gene expression RNA-seq data, which were batch-effects-normalized mRNA data (n=11,060), from the TCGA Pan-Cancer (PANCAN) dataset were downloaded from the University of California Santa Cruz Xena platform (http://xena.ucsc.edu). The PANCAN dataset included extracted cancer tissue data and normal tissue data for various cancer types, including stomach adenocarcinoma (STAD), esophageal carcinoma (ESCA), lung adenocarcinoma (LUAD), lung squamous cell carcinoma (LUSC), colon adenocarcinoma (COAD), breast invasive carcinoma (BRCA) and pancreatic adenocarcinoma (PAAD). Microarray datasets for gastric cancer, namely, GSE66229, GSE29272, and GSE27342, were obtained from the Gene Expression Omnibus database. The *PLA2G7* expression data were analyzed for the current study, and the log_2_(norm_value+1) values or the log_2_(norm_value) values for TCGA were used to represent the expression levels of the genes.

### In vivo Tumor xenograft studies

SNU-484 cells in the logarithmic growth phase were transplanted into immunodeficient nude mice (6 × 10^6^ cells in 200 μL of PBS + Matrigel/mouse). To determine the anticancer effect, the tumor size and weight of the mice were measured every 3 days. Tumor volume was calculated according to the formula tumor volume = longest diameter × shortest diameter × height/2. For intraperitoneal (i.p.) administration, the tumor was allowed to grow for approximately 2 weeks to reach an average volume of approximately 20 mm^3^. The mice were randomly divided into 4 groups: the vehicle control group (n = 7), the PACMA31 group (n = 5; 10 mg/kg), the darapladib group (n = 7; 10 mg/kg), the PACMA31 + darapladib group (n = 5) and the PACMA31 + darapladib group + fer-1 (10 mg/kg) (n = 5).

### Statistical analyses

All experiment was repeated at least three times. Data are presented as means ± SD (standard deviation). Statistical differences among groups were analyzed using GraphPad Prism 8 (GraphPad Software) and Excel. In all of the comparisons, a level of p < 0.05 was considered significant (n.s., not significant; *p < 0.05; **p < 0.01; ***p < 0.001).

## Data Availability

All study data are included in the article and supporting information

## Supplementary Figure Legends

**Supplementary Fig. 1. Metabolic library screening to identify a ferroptosis-targeting drug.**

(a) Heat map indicating the cell viability of 10 μM componuds alone and combination treatment with 0.5 μM RSL3. The colors indicate the relative cell viability levels. (b) Relative viability of Hs746T cells 0.5 μM RSL3 and/or several compounds in the presence or absence of Fer-1.

**Supplementary Fig. 2. Darapladib promotes RSL3-induced ferroptosis.**

(a) Relative viability of Hs746T and SNU-484 cells treated with various concentrations of darapladib for 20 h. (b) Relative viability of Hs746T and SNU-484 cells treated with 0.2 μM RSL3 and/or 2 μM darapladib in the presence or absence of 1 μM Fer-1, zVAD-fmk (z-VAD, 20 μM) or Necrostatin-1 (Nec-1, 30 μM). (c) LDH levels from Hs746T and SNU-484 cells treated with 0.2 μM RSL3 and/or 2 μM darapladib in the presence or absence of 1 μM Fer-1 for 20 h. (d) Relative viability of cells at high and low density upon various concentration of darapladib treatment in Hs746T and SNU-484 cells. Images of cells treated with indicated concentration of darapladib.

**Supplementary Fig. 3. Darapladib sensitizes various types of cancer cells to ferroptosis.**

(a) Relative viability of A549, H1299, HepG2 and YCC-16 cells treated with RSL3 or/and 2 μM darapladib in the presence or absence of Fer-1 for 20 h. (b) Relative viability of H9c2 and MEF cells treated with RSL3 or/and 2 μM darapladib (c) Relative viability of H9c2 cells treated with Erastin or/and 2 μM darapladib in the presence or absence of Fer-1 for 20 h.

**Supplementary Fig. 4. HDL re-sensitizes cells to ferroptosis under lipoprotein deficiency.**

(a) Western blot to determine whether lipoprotein deficiency affects the mTOR pathway in Hs746T and SNU-484 cells. (b) Relative viability of SNU-484 cells treated with RSL3 or/and 2 μM darapladib (B) Relative viability of SNU-484 cells treated with RSL3 and 2 μM darapladib cultured in medium containing FBS or LPDS for 6 h. Various lipoprotein were supplemented to medium containing LPDS.

**Supplementary Fig. 5. Lp-PLA2 does not contribute to the total levels of lysoPCs in the medium.**

(a) Levels of AA and lysoPC species in conditioned medium of Hs746T (1, 2, and 4 h) and SNU-484 (1 and 2 h) cells treated with 2 μM darapladib determined using LC–MS/MSThe intensities were normalized to the cell numbers.

**Supplementary Fig. 6. Knockdown of Lp-PLA2 augments RSL3-induced ferroptosis.**

(a) Analysis of Lp-PLA2 mRNA expression after 72 h of siRNA transfection in Hs746T and SNU-484 cells. (b) Relative viability of Hs746T and SNU-484 cells treated with RSL3 after 72 h of siRNA transfection. The relative expression levels were normalized to the β-actin expression levels.

**Supplementary Fig. 7. Lp-PLA2 negatively regulates lipid peroxidation and ferroptosis.**

(a) Analysis of mRNA expression in Lp-PLA2 depletion cells. Relative expression levels were normalized to the β-actin expression levels. (b and c) Relative cell viability and LDH levels in WT and *PLA2G7* KO YCC-16 cells treated with RSL3 for 72 h. (d) Crystal violet staining result after various concentration of RSL3 treatment in WT and *PLA2G7* KO YCC-16 cells. (e and f) Relative cell viability and LDH levels in WT and *PLA2G7* KO YCC-16 cells treated with RSL3 in the presence or absence of Fer-1 for 72 h. (g) Lipid peroxidation level in WT and *PLA2G7* KO YCC-16 cells treated with RSL3 for 2 h. (h) Cell viability from WT and *PLA2G7* KO YCC-16 cells cultured with cysteine deficient medium for 48 h. (i) Western blots showing the expression levels of well-known ferroptosis regulators, such as GPX4, NRF2, FSP1, ACSL4, LPCAT3, PEBP1, FADS1, and ELOVL5, in Lp-PLA2-depleted H1299 and YCC-16 cells. (j) Relative cell viability from WT and *PLA2G7* KO YCC-16 cells treated with 4 µM RSL3 or/and 0.5 µM darapladib for 48 h. (k) Relative viability of Hs746T cells treated with various concentration of RSL3 after ectopic expression of Lp-PLA2. (l) Lipid peroxidation level in Hs746T cells treated with various concentration of RSL3 after ectopic expression of Lp-PLA2.

**Supplementary Fig. 8. PE and PE-p species accumulate, while lysoPE and lysoPC species are downregulated in *Lp-PLA2* KO H1299 cells.**

(a) Proportions of various lipid classes in WT and *PLA2G7* KO H1299 cells were detected using LC–MS/MS. (b) Levels of representative lipid species in WT and *PLA2G7* KO H1299 cells detected using LC–MS/MS. The intensities were normalized to the total sum of the peak areas. The data are the means ± SDs (n=8 independent experiments). (c) Lp-PLA2 downregulates genes involved in de novo fatty acid synthesis. Western blots showing the expression levels of well-known fatty acid metabolism-related genes, such as FASN, ACC, ACC-p and SREBP1 in WT and *PLA2G7* KO H1299 cells

**Supplementary Fig. 9. PE and PE-p species accumulate, while lysoPE, lysoPE-p, and MUFA species are downregulated, in response to darapladib.**

(a) Proportions of various lipid classes in Hs746T cells treated with darapladib. (b) Volcano plot of lipid classes showing the log2(fold change) and –log10(p) values in control vs. darapladib-treated Hs746T cells. (c) Levels of representative lipid species in Hs746T cells treated with darapladib for 1, 2, and 4 h were detected using LC– MS/MS. The intensities were normalized to the total sum of the peak areas. The data are the means ± SDs (n=7 independent experiments). (d) Darapladib downregulates genes involved in de novo fatty acid synthesis. Western blots showing the expression levels of well-known fatty acid metabolism-related genes, such as FASN, ACC, ACC-p, PPAR gamma, and SREBP1, after treatment with darapladib for the indicated times.

**Supplementary Fig. 10. Oxidized PE-C18:0/C20:4 and PE-C18:0/C20:4 are up-regulated, in response to darapladib and in *PLA2G7* KO cells**

(a) Concentrations of oxidized PE-C18:0/C20:4, PE-C18:0/C20:4, and PE-C18:0/C22:4 after treatment with RSL3 or/and 2 μM darapladib for 3 h in Hs746T cells. The ratios of oxidized PE-C18:0/C20:4 to PE-C18:0/C20:4 are also shown. The levels of oxidized PE, PE-C18:0/C20:4, and PE-C18:0/22:4 were determined using LC– MS/MS. The concentrations were normalized to the cellular protein level. The data are the means ± SDs (n=6 independent experiments). (b) Concentrations of oxidized PE-C18:0/C20:4, PE-C18:0/C20:4, and PE-C18:0/C22:4 in WT and *PLA2G7* KO H1299 cells. The ratios of oxidized PE-C18:0/C20:4 to PE-C18:0/C20:4 are also shown. The levels of oxidized PE and PE-C18:0/C20:4 were determined using LC–MS/MS. The concentrations were normalized to the cellular protein level. The data are the means ± SDs (n=5 independent experiments).

**Supplementary Fig. 11. Expression of *PLA2G7* in several types of tumors and normal tissues from transcriptome datasets.**

(a) Upregulation of *PLA2G7* in several cancer types, including gastric cancer. Expression of *PLA2G7* in stomach adenocarcinoma (STAD), esophageal carcinoma (ESCA), lung adenocarcinoma (LUAD), lung squamous cell carcinoma (LUSC), colon adenocarcinoma (COAD), breast invasive carcinoma (BRCA) and pancreatic adenocarcinoma (PAAD) tissues and their matched normal tissues. The expression data were extracted from the TCGA repository. (b) Relative expression levels of *PLA2G7* in gastric cancer and normal tissues from independent transcriptome datasets (GSE66229, GSE29272, and GSE27342).

**Supplementary Fig. 12. GPX4 deficient cell more sensitive to ferroptosis in response to darapladib.**

(A) Western blot confirming the expression of GPX4 protein in GPX4 depletion cells. (B) Images of GPX4 depleted cells treated with 5 μM darapladib and/or 2 μM fer-1. (C) Relative cell viability in WT and GPX4 depleted H1299 cells treated with 5 μM darapladib in the presence or absence of 2 μM Fer-1 for 20 h.

**Supplementary Fig. 13. PACMA31 is a reliable ferroptosis inducers.**

(a) Relative viability of Hs746T, SNU-484 and H1299 cells treated with various concentration of PACMA31. (b) Relative viability of Hs746T, SNU-484 and H1299 cells treated with PACMA31 in the presence or absence of Fer-1 for 20 h. (c) Lipidomic analysis acquired from the UPLC/QTOF MS spectra in xenografted tumor treated with PACMA31 and/or darapladib in the presence or absence of Fer-1. Each value in the heatmap is a colored representation of a calculated Z score. Lipids are presented as the total number of carbon atoms and the total number of double bonds. n = 10 independent experiments.

**Supplementary Fig. 14. Lp-PLA2 is a common regulator of ferroptosis.**

(a) Relative viability of SNU-484 and H1299 cells treated with RSL3 following a 72 h transfection of siRNA against the genes encoding sPLA2, cPLA2, iPLA2, and Lp-PLA2 (*PLA2G2A, PLA2G4A, PLA2G6*, and *PLA2G7*). (b and c) Relative viability of Hs746T and H9c2 cells treated with several inhibitors for PLA2, such as (S)-BEL (an iPLA2 inhibitor), MAFP (a cPLA2/sPLA2 inhibitor), MJ33 (an inhibitor of PLA2 activity of PRDX6) or/and RSL3 in the presence or absence of Fer-1 for 20 h.

**Supplementary Fig. 15. Supplementation with lysoPC18:0 or lysoPC16:0 abrogates RSL3-induced ferroptosis.**

(a) Relative viability of Hs746T cells pretreated with lysoPC (16:0 or 18:1) for 1 h and treated with RSL3 or/and 2 μM darapladib for 20 h.

## Acknowledgments

This work was supported by a grant from the KRIBB Research Initiative Program and Korea Basic Science Institute (C270000) and by National Research Foundation of Korea (NRF) grants funded by the Korean government (MSIT) (Nos. 2017M3A9G5083321, 2019M3A9D5A01102796, 2020R1A2C1006841, 2020R1A2C2007835, and 2022R1A2C4002108).

## Author Contributions

M.O., S.Y.J., J.-Y.L., S.C.L., B.-S.H., G.-S.H., and E.-W.L. conceived and designed the study and wrote the manuscript; M.O., J.-Y.L., and J.W.K. performed most of the biochemical experiments; S.Y.J., Y.J., D.K., and G.-S.H. performed the lipidomics experiments; M.O., J.W.K. J.S., and T.-S.H. performed the mouse xenograft experiments; T.-S.H., E.J., and H.Y.S. analyzed gene expression from TCGA and the other databases; and M.W.K., K.-H.S., K.-J.O., W.-K.K., K.-H.B., and Y.-M.H. analyzed the data and provided useful comments.

## Declaration of Interests

The authors declare no competing interests.

## Notes

### Competing Interest Statement

The authors have declared no competing interest.

