## Supplement fig for "Darapladib, an inhibitor of Lp-PLA2, sensitizes cancer cells to ferroptosis by remodeling lipid metabolism"

**Figure S1**

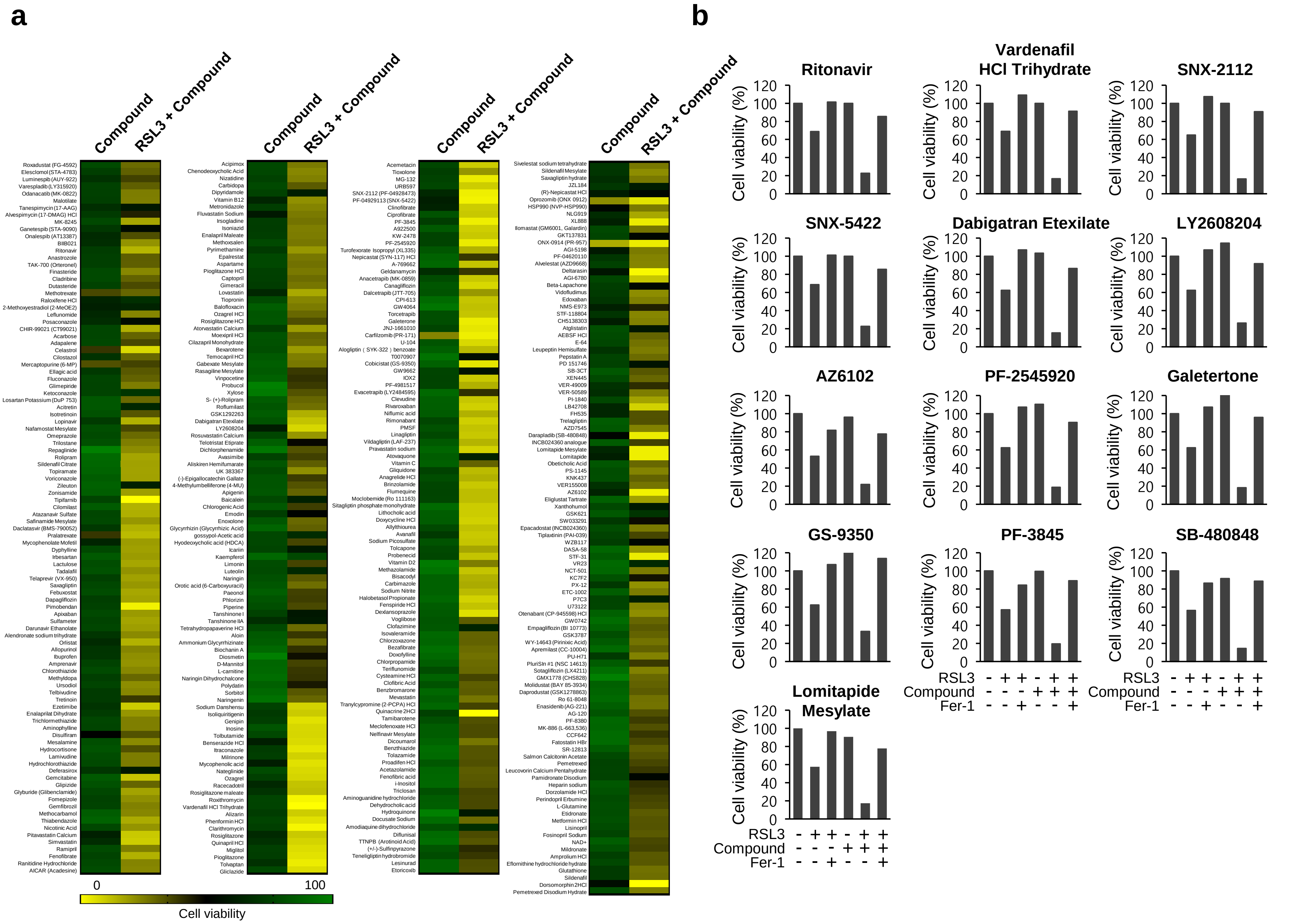

a
Compound
RSL3 + Compound
Roxadustat (FG-4592)
Elesclomol (STA-4783)
Luminespib (AUY-922)
Varespladib (LY315920)
Odanacatib (MK-0822)
Malotilate
Tanespimycin (17-AAG)
Alvespimycin (17-DMAG) HCl
MK-8245
Ganetespib (STA-9090)
Onalespib (AT13387)
BIIB021
Ritonavir
Anastrozole
TAK-700 (Orteronel)
Finasteride
Cladribine
Dutasteride
Methotrexate
Raloxifene HCl
2-Methoxyestradiol (2-MeOE2)
Leflunomide
Posaconazole
CHIR-99021 (CT99021)
Acarbose
Adapalene
Celastrol
Cilostazol
Mercaptopurine (6-MP)
Ellagic acid
Fluconazole
Glimepiride
Ketoconazole
Losartan Potassium (DuP 753)
Acitretin
Isotretinoin
Lopinavir
Nafamostat Mesylate
Omeprazole
Trilostane
Repaglinide
Rolipram
Sildenafil Citrate
Topiramate
Voriconazole
Zileuton
Zonisamide
Tipifarnib
Cilomilast
Atazanavir Sulfate
Safinamide Mesylate
Daclatasvir (BMS-790052)
Pralatrexate
Mycophenolate Mofetil
Dyphylline
Irbesartan
Lactulose
Tadalafil
Telaprevir (VX-950)
Saxagliptin
Febuxostat
Dapagliflozin
Pimobendan
Apixaban
Sulfameter
Darunavir Ethanolate
Alendronate sodium trihydrate
Orlistat
Allopurinol
Ibuprofen
Amprenavir
Chlorothiazide
Methyldopa
Ursodiol
Telbivudine
Tretinoin
Ezetimibe
Enalaprilat Dihydrate
Trichlormethiazide
Aminophylline
Disulfiram
Mesalamine
Hydrocortisone
Lamivudine
Hydrochlorothiazide
Deferasirox
Gemcitabine
Glipizide
Glyburide (Glibenclamide)
Fomepizole
Gemfibrozil
Methocarbamol
Thiabendazole
Nicotinic Acid
Pitavastatin Calcium
Simvastatin
Ramipril
Fenofibrate
Ranitidine Hydrochloride
AICAR (Acadesine)
Acipimox
Chenodeoxycholic Acid
Nizatidine
Carbidopa
Dipyridamole
Vitamin B12
Metronidazole
Fluvastatin Sodium
Irsogladine
Isoniazid
Enalapril Maleate
Methoxsalen
Pyrimethamine
Epalrestat
Aspartame
Pioglitazone HCl
Captopril
Gimeracil
Lovastatin
Tiopronin
Balofloxacin
Ozagrel HCl
Rosiglitazone HCl
Atorvastatin Calcium
Moexipril HCl
Cilazapril Monohydrate
Bexarotene
Temocapril HCl
Gabexate Mesylate
Rasagiline Mesylate
Vinpocetine
Probucol
Xylose
S- (+)-Rolipram
Roflumilast
GSK1292263
Dabigatran Etexilate
LY2608204
Rosuvastatin Calcium
Telotristat Etiprate
Dichlorphenamide
Avasimibe
Aliskiren Hemifumarate
UK 383367
(-)-Epigallocatechin Gallate
4-Methylumbelliferone (4-MU)
Apigenin
Baicalein
Chlorogenic Acid
Emodin
Enoxolone
Glycyrrhizin (Glycyrrhizic Acid)
gossypol-Acetic acid
Hyodeoxycholic acid (HDCA)
Icariin
Kaempferol
Limonin
Luteolin
Naringin
Orotic acid (6-Carboxyuracil)
Paeonol
Phlorizin
Piperine
Tanshinone I
Tanshinone IIA
Tetrahydropapaverine HCl
Aloin
Ammonium Glycyrrhizinate
Biochanin A
Diosmetin
D-Mannitol
L-carnitine
Naringin Dihydrochalcone
Polydatin
Naringenin
Sodium Danshensu
Isoliquiritigenin
Genipin
Inosine
Tolbutamide
Benserazide HCl
Itraconazole
Milrinone
Mycophenolic acid
Nateglinide
Ozagrel
Racecadotril
Rosiglitazone maleate
Roxithromycin
Vardenafil HCl Trihydrate
Alizarin
Phenformin HCl
Clarithromycin
Rosiglitazone
Quinapril HCl
Miglitol
Pioglitazone
Tolvaptan
Gliclazide
Acemetacin
Tioxolone
MG-132
URB597
SNX-2112 (PF-04928473)
PF-04929113 (SNX-5422)
Clinofibrate
Ciprofibrate
PF-3845
A922500
KW-2478
PF-2545920
Turofexorate Isopropyl (XL335)
Nepicastat (SYN-117) HCl
A-769662
Geldanamycin
Anacetrapib (MK-0859)
Canagliflozin
Dalcetrapib (JTT-705)
CPI-613
GW4064
Torcetrapib
Galeterone
JNJ-1661010
Carfilzomib (PR-171)
U-104
Alogliptin ( SYK-322 ) benzoate
T0070907
Cobicistat (GS-9350)
GW9662
IOX2
PF-4981517
Evacetrapib (LY2484595)
Clevudine
Rivaroxaban
Niflumic acid
Rimonabant
PMSF
Linagliptin
Vildagliptin (LAF-237)
Pravastatin sodium
Atovaquone
Vitamin C
Gliquidone
Anagrelide HCl
Brinzolamide
Flumequine
Moclobemide (Ro 111163)
Sitagliptin phosphate monohydrate
Lithocholic acid
Doxycycline HCl
Allylthiourea
Avanafil
Sodium Picosulfate
Tolcapone
Probenecid
Vitamin D2
Methazolamide
Bisacodyl
Carbimazole
Sodium Nitrite
Halobetasol Propionate
Fenspiride HCl
Dexlansoprazole
Voglibose
Clofazimine
Isovaleramide
Chlorzoxazone
Bezafibrate
Doxofylline
Chlorpropamide
Teriflunomide
Cysteamine HCl
Clofibric Acid
Benzbromarone
Mevastatin
Tranylcypromine (2-PCPA) HCl
Quinacrine 2HCl
Tamibarotene
Meclofenoxate HCl
Nelfinavir Mesylate
Dicoumarol
Benzthiazide
Tolazamide
Proadifen HCl
Acetazolamide
Fenofibric acid
i-Inositol
Triclosan
Aminoguanidine hydrochloride
Dehydrocholic acid
Hydroquinone
Docusate Sodium
Amodiaquine dihydrochloride
Diflunisal
TTNPB (Arotinoid Acid)
(+/-)-Sulfinpyrazone
Teneligliptin hydrobromide
Lesinurad
Etoricoxib
Sivelestat sodium tetrahydrate
Sildenafil Mesylate
Saxagliptin hydrate
JZL184
(R)-Nepicastat HCl
Oprozomib (ONX 0912)
HSP990 (NVP-HSP990)
NLG919
XL888
Ilomastat (GM6001, Galardin)
GKT137831
ONX-0914 (PR-957)
AGI-5198
PF-04620110
Alvelestat (AZD9668)
Deltarasin
AGI-6780
Beta-Lapachone
Vidofludimus
Edoxaban
NMS-E973
STF-118804
CH5138303
Atglistatin
AEBSF HCl
E-64
Leupeptin Hemisulfate
Pepstatin A
PD 151746
SB-3CT
XEN445
VER-49009
VER-50589
PI-1840
LB42708
FH535
Trelagliptin
AZD7545
Darapladib (SB-480848)
INCB024360 analogue
Lomitapide Mesylate
Lomitapide
Obeticholic Acid
PS-1145
KNK437
VER155008
AZ6102
Eliglustat Tartrate
Xanthohumol
GSK621
SW033291
Epacadostat (INCB024360)
Tiplaxtinin (PAI-039)
WZB117
DASA-58
STF-31
VR23
NCT-501
KC7F2
PX-12
ETC-1002
P7C3
U73122
Otenabant (CP-945598) HCl
GW0742
Empagliflozin (BI 10773)
GSK3787
WY-14643 (Pirinixic Acid)
Apremilast (CC-10004)
PU-H71
PluriSIn #1 (NSC 14613)
Sotagliflozin (LX4211)
GMX1778 (CHS828)
Molidustat (BAY 85-3934)
Daprodustat (GSK1278863)
Ro 61-8048
Enasidenib (AG-221)
AG-120
PF-8380
MK-886 (L-663,536)
CCF642
Fatostatin HBr
SR-12813
Salmon Calcitonin Acetate
Leucovorin Calcium Pentahydrate
Pemetrexed
Pamidronate Disodium
Heparin sodium
Dorzolamide HCl
Perindopril Erbumine
L-Glutamine
Etidronate
Metformin HCl
Lisinopril
Fosinopril Sodium
NAD+
Mildronate
Amprolium HCl
Eflornithine hydrochloride hydrate
Glutathione
Sildenafil
Dorsomorphin 2HCl
Pemetrexed Disodium Hydrate
0
100
Cell viability
b
Ritonavir
Vardenafil HCl Trihydrate
SNX-2112
SNX-5422
Dabigatran Etexilate
LY2608204
AZ6102
PF-2545920
Galetertone
GS-9350
PF-3845
SB-480848
Lomitapide Mesylate
Cell viability (%)
0
20
40
60
80
100
120
RSL3 - + + - + +
Compound - - - + + +
Fer-1 - - + - - +

### Figure S2

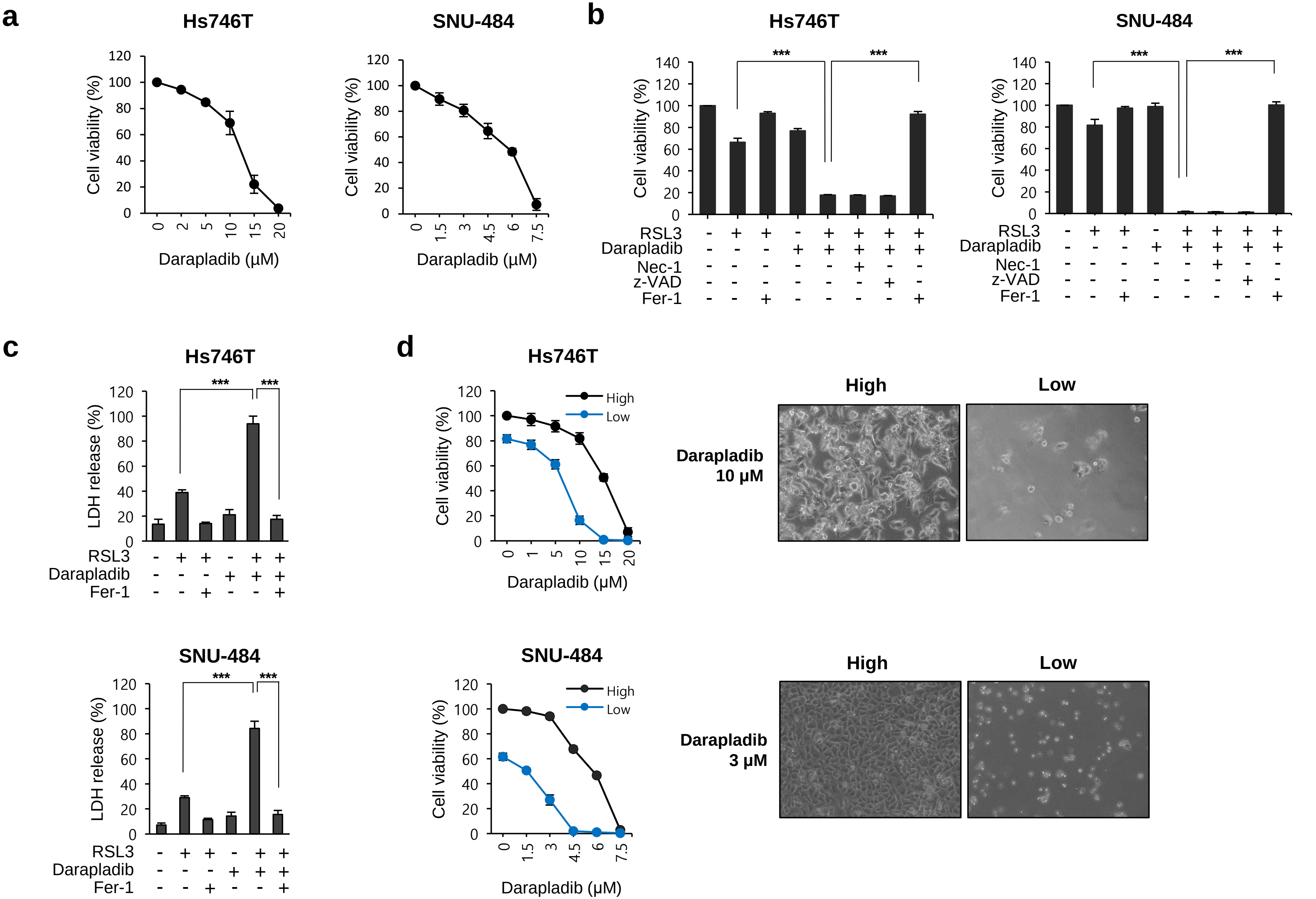
a
Hs746T
SNU-484
Cell viability (%)
Darapladib (μM)
b
Hs746T
SNU-484
***
RSL3
Darapladib
Nec-1
z-VAD
Fer-1
c
Hs746T
SNU-484
LDH release (%)
d
High
Low
Darapladib 10 μM
Darapladib 3 μM

### Figure S3

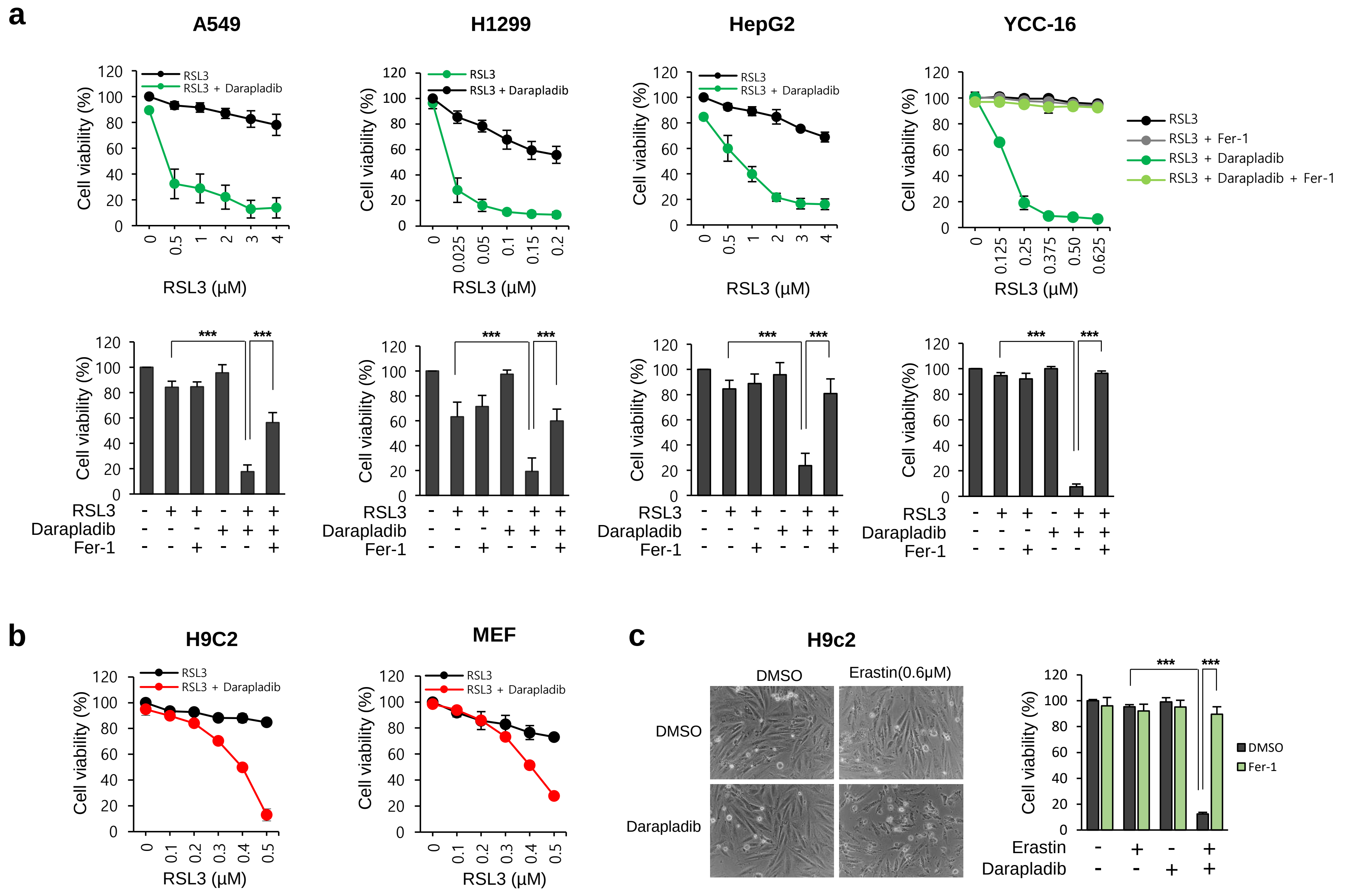
a
A549
H1299
HepG2
YCC-16
Cell viability (%)
RSL3
RSL3 + Darapladib
RSL3 + Fer-1
RSL3 + Darapladib + Fer-1
RSL3 (μM)
***
RSL3
Darapladib
Fer-1
b
H9C2
MEF
c
H9c2
DMSO
Erastin(0.6μM)
Darapladib
Fer-1
Erastin

Figure S4

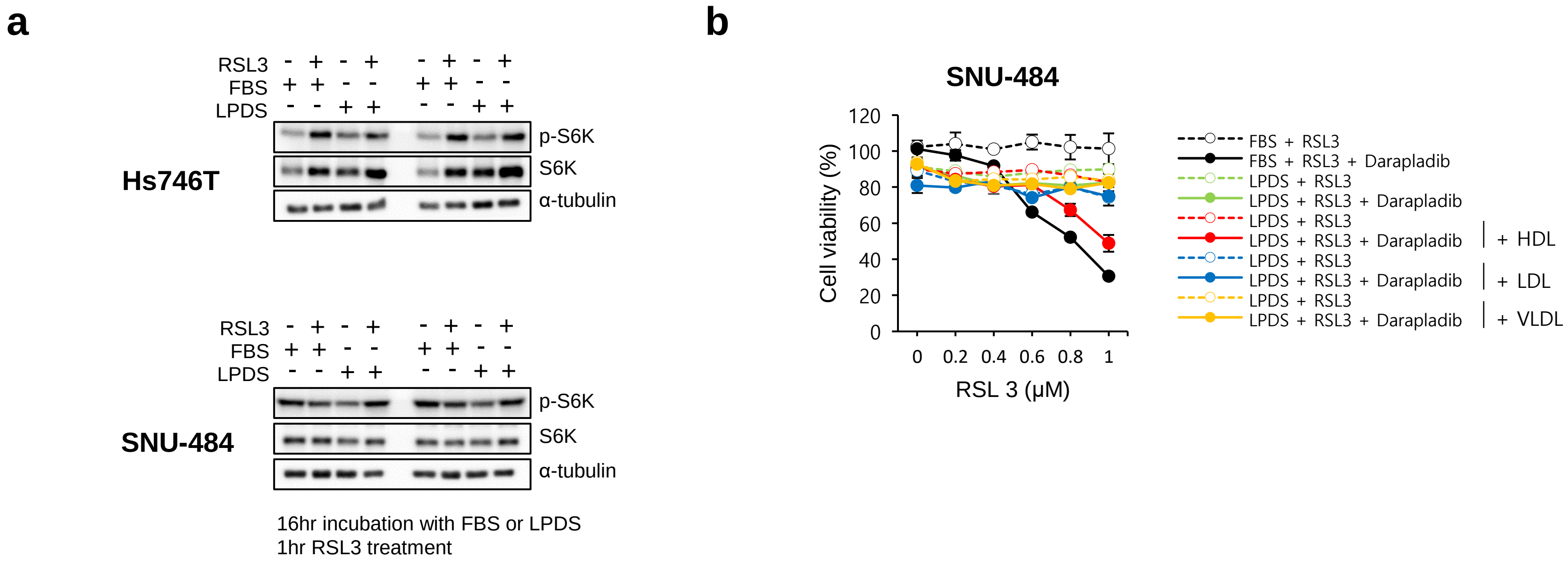
a
RSL3 - + - + - + - +
FBS + + - - + + - -
LPDS - - + + - - + +
Hs746T
p-S6K
S6K
α-tubulin
RSL3 - + - + - + - +
FBS + + - - + + - -
LPDS - - + + - - + +
SNU-484
p-S6K
S6K
α-tubulin
16hr incubation with FBS or LPDS
1hr RSL3 treatment
b
SNU-484
Cell viability (%)
120
100
80
60
40
20
0
0 0.2 0.4 0.6 0.8 1
RSL 3 (μM)
FBS + RSL3
FBS + RSL3 + Darapladib
LPDS + RSL3
LPDS + RSL3 + Darapladib
LPDS + RSL3
LPDS + RSL3 + Darapladib
+ HDL
LPDS + RSL3
LPDS + RSL3 + Darapladib
+ LDL
LPDS + RSL3
LPDS + RSL3 + Darapladib
+ VLDL

Figure S5

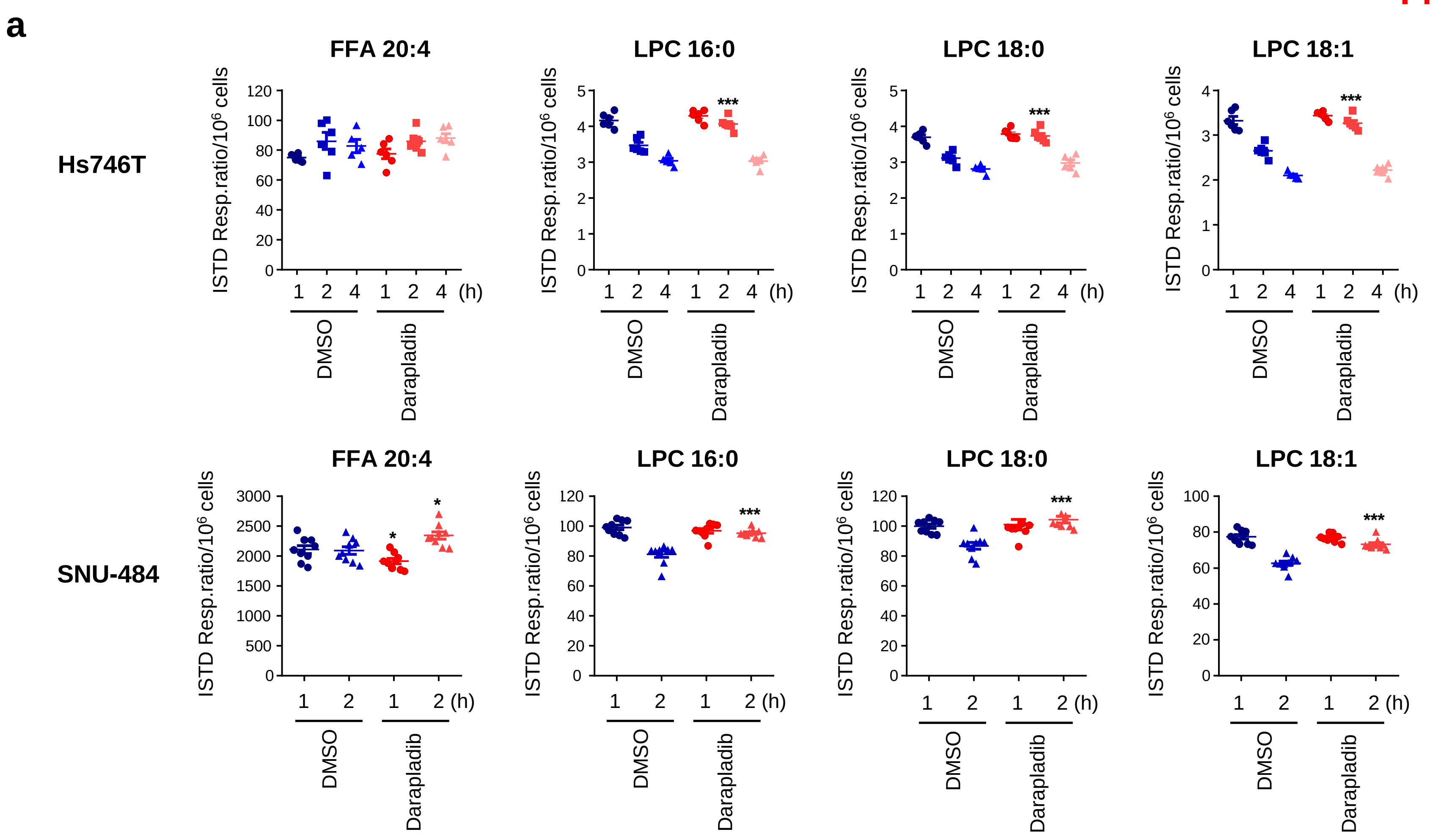
a
Hs746T
SNU-484
FFA 20:4
LPC 16:0
LPC 18:0
LPC 18:1
ISTD Resp.ratio/10^6 cells
***
*
1 2 4 (h)
1 2 (h)
DMSO
Darapladib

Figure S6

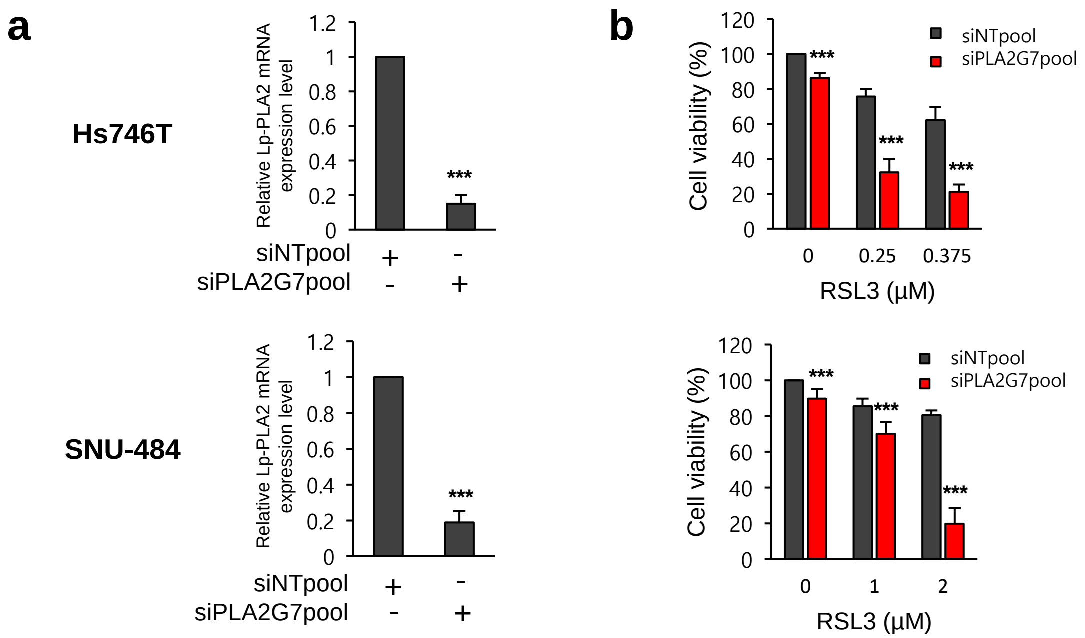
a
b
Hs746T
SNU-484
Relative Lp-PLA2 mRNA expression level
1.2
1
0.8
0.6
0.4
0.2
0
***
siNTpool + -
siPLA2G7pool - +
Cell viability (%)
120
100
80
60
40
20
0
siNTpool
siPLA2G7pool
0 0.25 0.375
RSL3 (μM)
0 1 2
RSL3 (μM)

### Figure S7

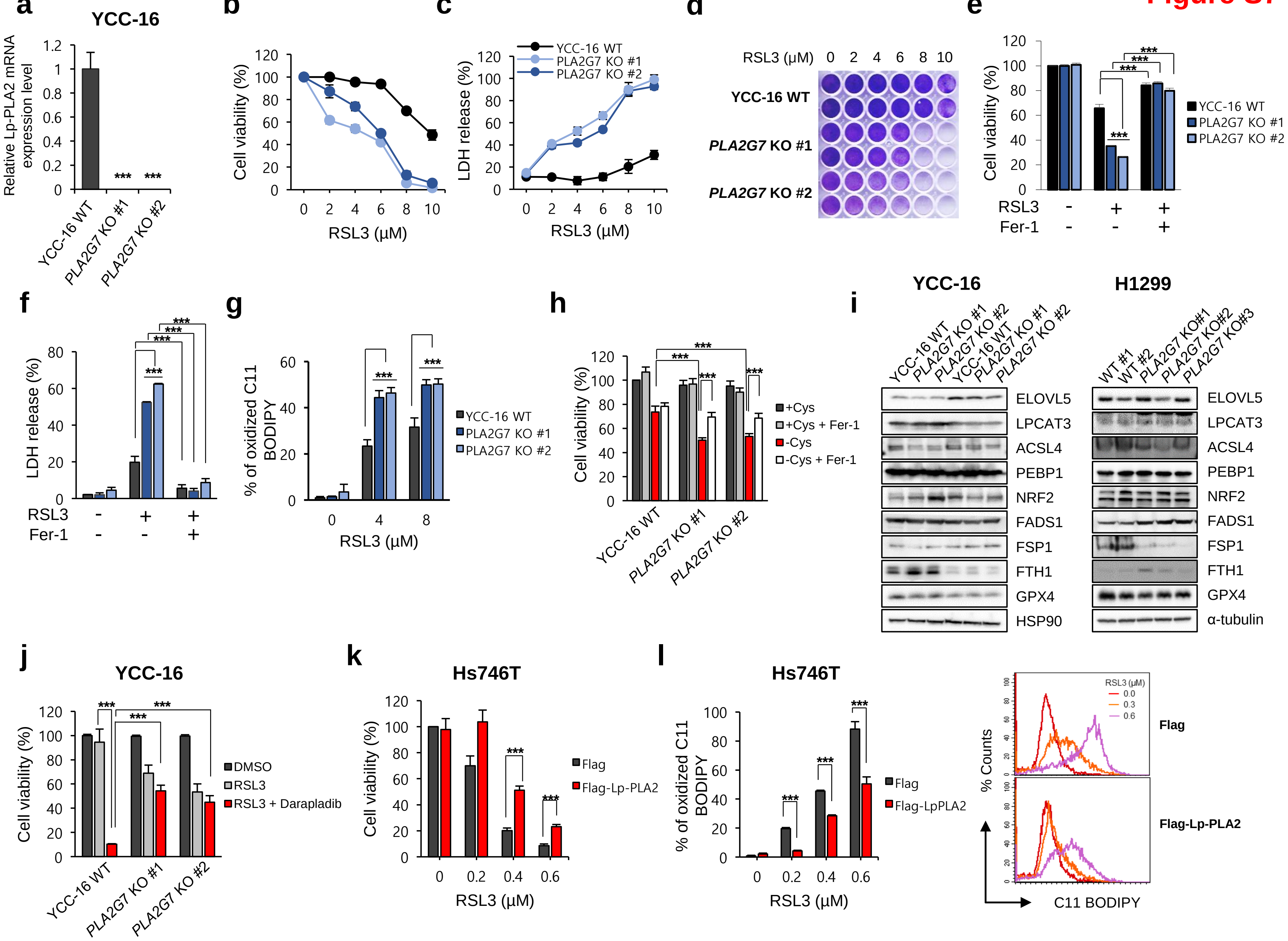
a
YCC-16
Relative Lp-PLA2 mRNA expression level
YCC-16 WT
PLA2G7 KO #1
PLA2G7 KO #2
***
b
Cell viability (%)
RSL3 (μM)
c
LDH release (%)
YCC-16 WT
PLA2G7 KO #1
PLA2G7 KO #2
d
RSL3 (μM) 0 2 4 6 8 10
YCC-16 WT
PLA2G7 KO #1
PLA2G7 KO #2
e
RSL3
Fer-1
f
g
% of oxidized C11 BODIPY
h
+Cys
+Cys + Fer-1
-Cys
-Cys + Fer-1
i
YCC-16
H1299
WT #1
WT #2
PLA2G7 KO#1
PLA2G7 KO#2
PLA2G7 KO#3
ELOVL5
LPCAT3
ACSL4
PEBP1
NRF2
FADS1
FSP1
FTH1
GPX4
HSP90
α-tubulin
j
DMSO
RSL3
RSL3 + Darapladib
k
Hs746T
Flag
Flag-Lp-PLA2
l
Flag-LpPLA2
% Counts
C11 BODIPY
RSL3 (μM)
0.0
0.3
0.6

### Figure S8

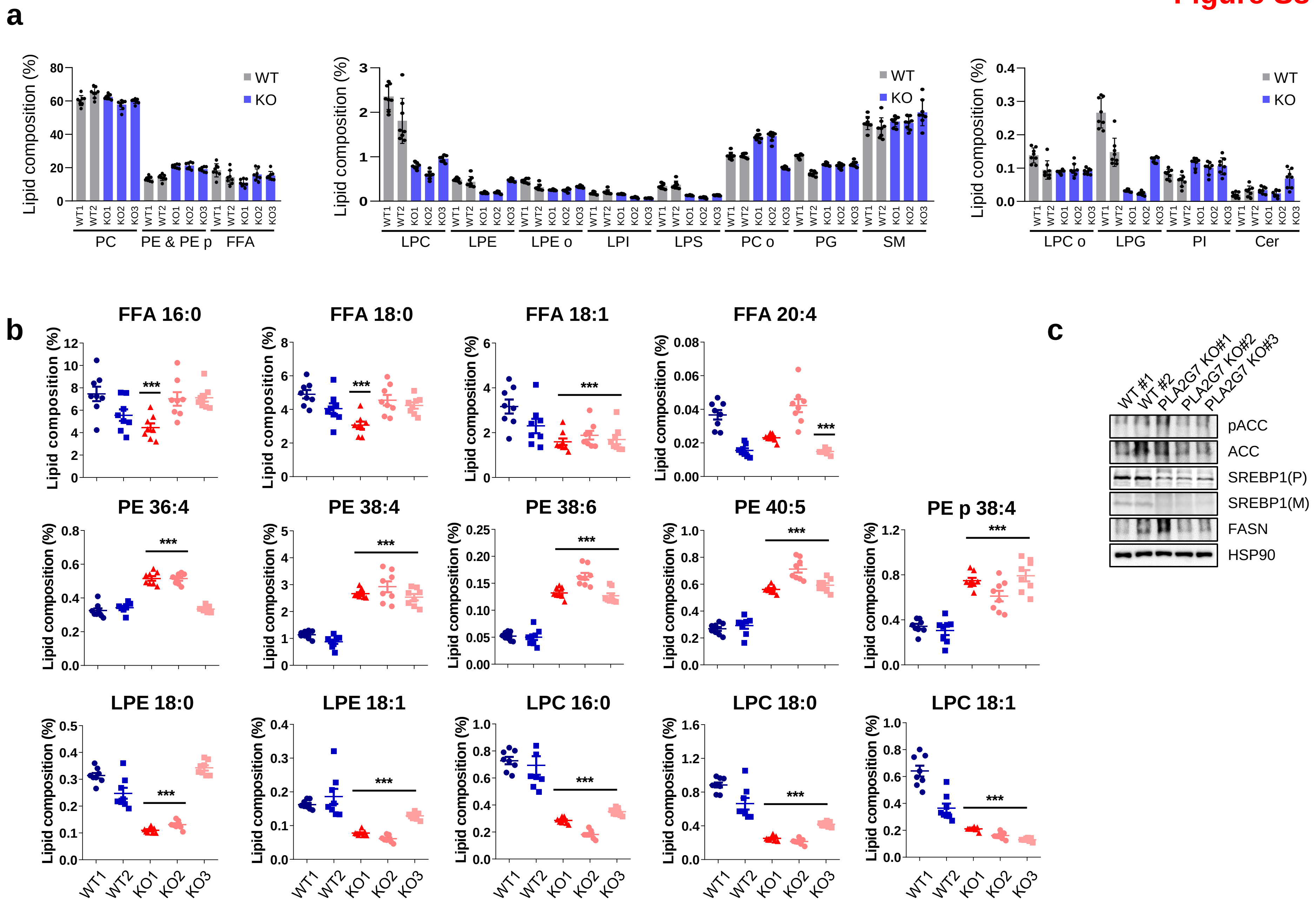

a
Lipid composition (%)
WT
KO
WT1 WT2 KO1 KO2 KO3
PC
PE & PE p
FFA
LPC
LPE
LPE o
LPI
LPS
PC o
PG
SM
LPC o
LPG
PI
Cer
b
FFA 16:0
FFA 18:0
FFA 18:1
FFA 20:4
PE 36:4
PE 38:4
PE 38:6
PE 40:5
PE p 38:4
LPE 18:0
LPE 18:1
LPC 16:0
LPC 18:0
LPC 18:1
***
c
WT #1
WT #2
PLA2G7 KO#1
PLA2G7 KO#2
PLA2G7 KO#3
pACC
ACC
SREBP1(P)
SREBP1(M)
FASN
HSP90

### Figure S9

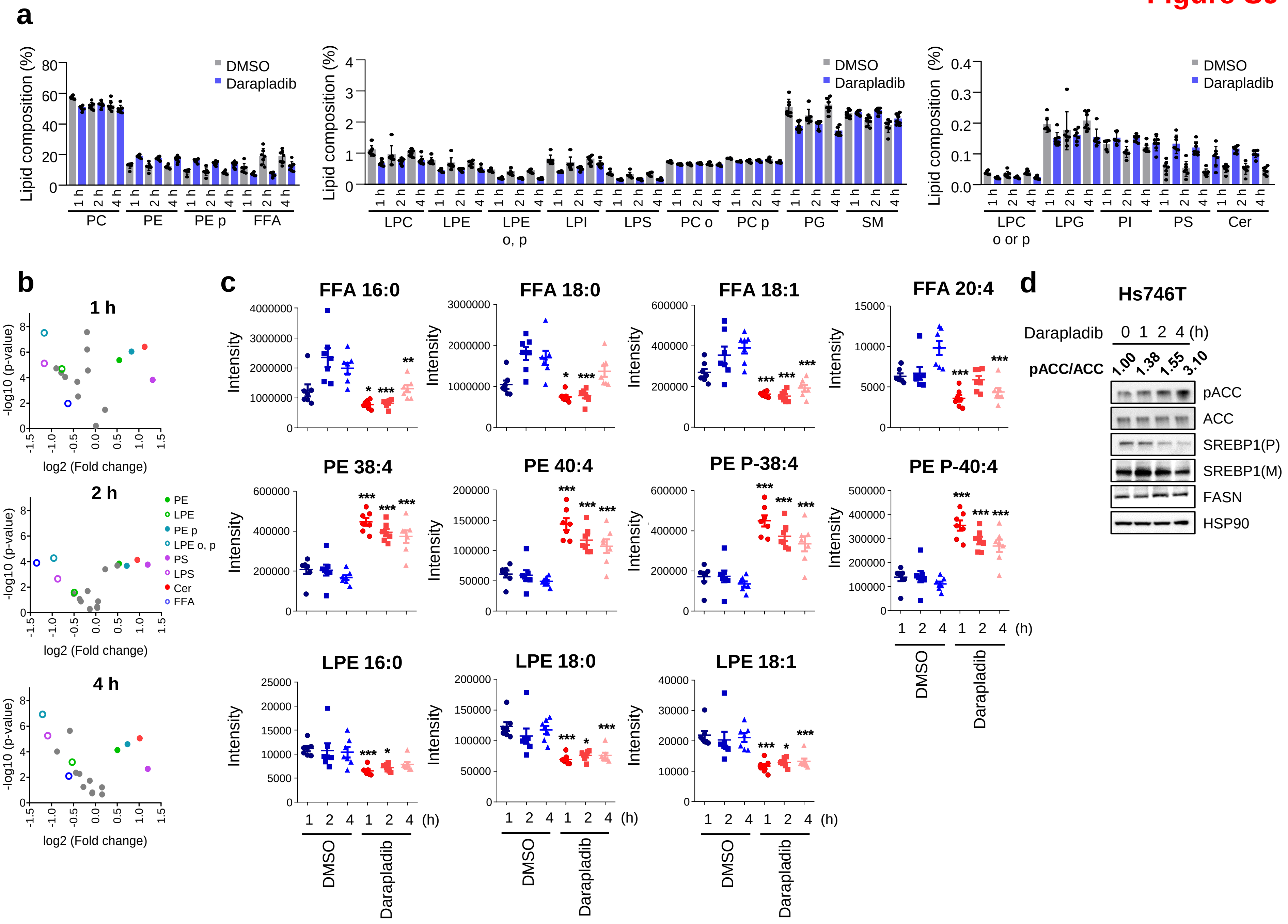
a
Lipid composition (%)
DMSO
Darapladib
1 h
2 h
4 h
PC
PE
PE p
FFA
LPC
LPE
LPE o, p
LPI
LPS
PC o
PC p
PG
SM
LPC o or p
LPG
PI
PS
Cer
b
1 h
2 h
4 h
-log10 (p-value)
log2 (Fold change)
PE
LPE
PE p
LPE o, p
PS
LPS
Cer
FFA
c
FFA 16:0
FFA 18:0
FFA 18:1
FFA 20:4
PE 38:4
PE 40:4
PE P-38:4
PE P-40:4
LPE 16:0
LPE 18:0
LPE 18:1
Intensity
(h)
DMSO
Darapladib
d
Hs746T
Darapladib 0 1 2 4 (h)
pACC/ACC 1.00 1.38 1.55 3.10
pACC
ACC
SREBP1(P)
SREBP1(M)
FASN
HSP90

Fig. S10

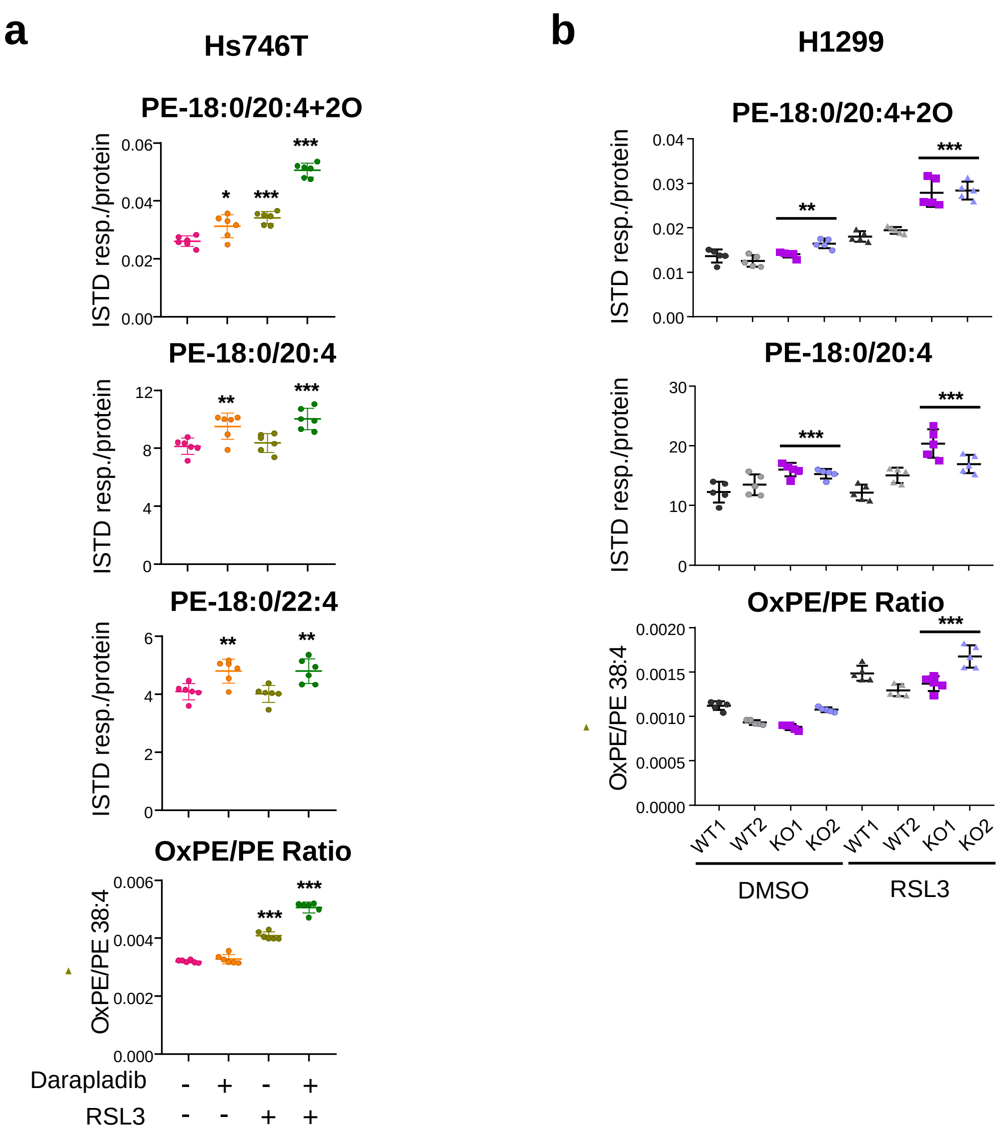

a
Hs746T
PE-18:0/20:4+2O
ISTD resp./protein
0.06
0.04
0.02
0.00
*
***
***
PE-18:0/20:4
12
8
4
0
**
***
PE-18:0/22:4
6
4
2
0
**
**
OxPE/PE Ratio
OxPE/PE 38:4
0.006
0.004
0.002
0.000
***
***
Darapladib - + - +
RSL3 - - + +
b
H1299
PE-18:0/20:4+2O
ISTD resp./protein
0.04
0.03
0.02
0.01
0.00
**
***
PE-18:0/20:4
30
20
10
0
***
***
OxPE/PE Ratio
OxPE/PE 38:4
0.0020
0.0015
0.0010
0.0005
0.0000
***
WT1 WT2 KO1 KO2 WT1 WT2 KO1 KO2
DMSO
RSL3

**Figure S11**

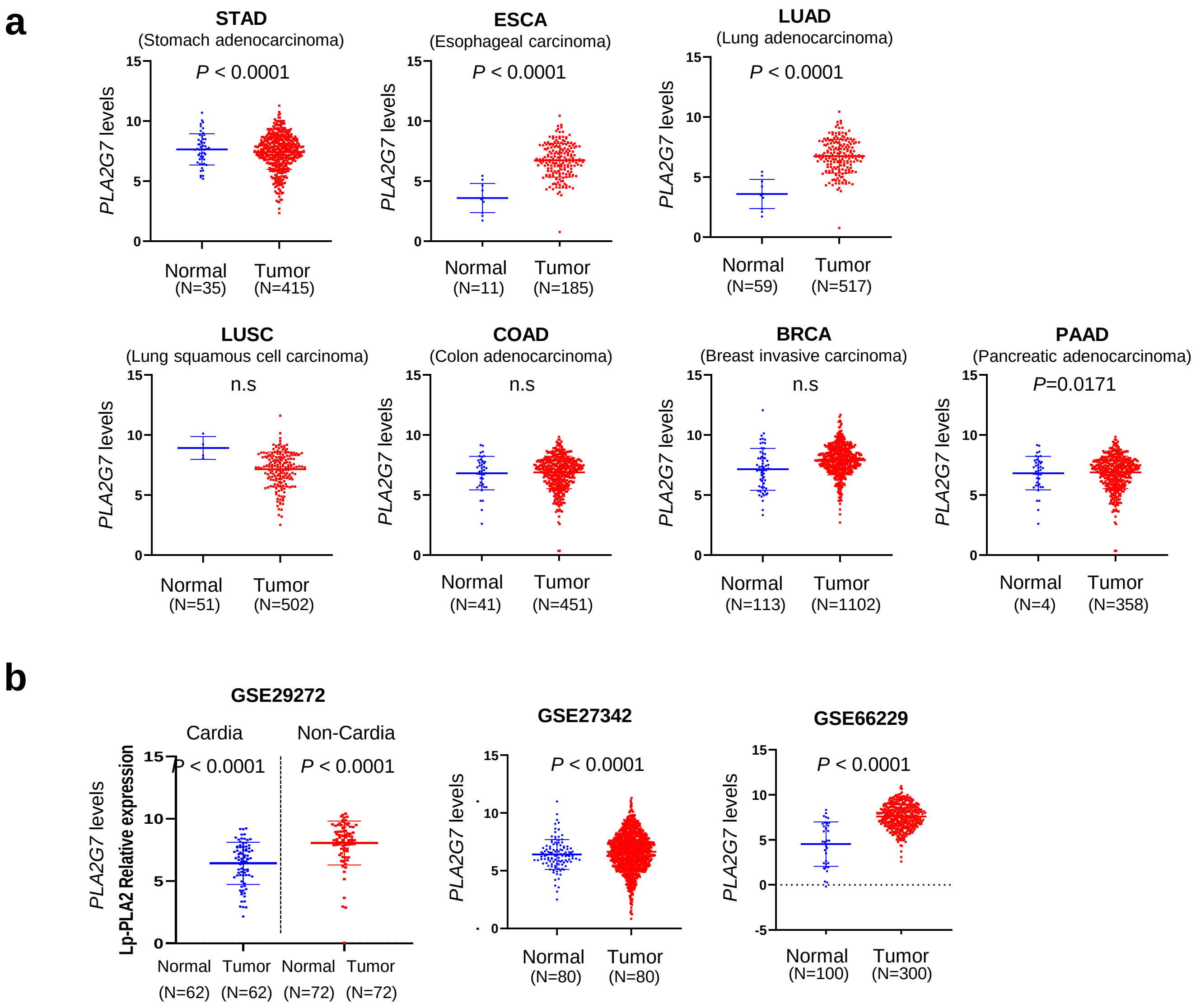
a
STAD
(Stomach adenocarcinoma)
P < 0.0001
PLA2G7 levels
Normal (N=35)
Tumor (N=415)
ESCA
(Esophageal carcinoma)
P < 0.0001
Normal (N=11)
Tumor (N=185)
LUAD
(Lung adenocarcinoma)
P < 0.0001
Normal (N=59)
Tumor (N=517)
LUSC
(Lung squamous cell carcinoma)
n.s
Normal (N=51)
Tumor (N=502)
COAD
(Colon adenocarcinoma)
n.s
Normal (N=41)
Tumor (N=451)
BRCA
(Breast invasive carcinoma)
n.s
Normal (N=113)
Tumor (N=1102)
PAAD
(Pancreatic adenocarcinoma)
P=0.0171
Normal (N=4)
Tumor (N=358)
b
GSE29272
Cardia
Non-Cardia
P < 0.0001
P < 0.0001
PLA2G7 levels
Lp-PLA2 Relative expression
Normal (N=62)
Tumor (N=62)
Normal (N=72)
Tumor (N=72)
GSE27342
P < 0.0001
Normal (N=80)
Tumor (N=80)
GSE66229
P < 0.0001
Normal (N=100)
Tumor (N=300)

Figure S12

a

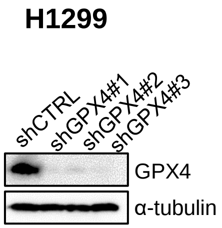


b

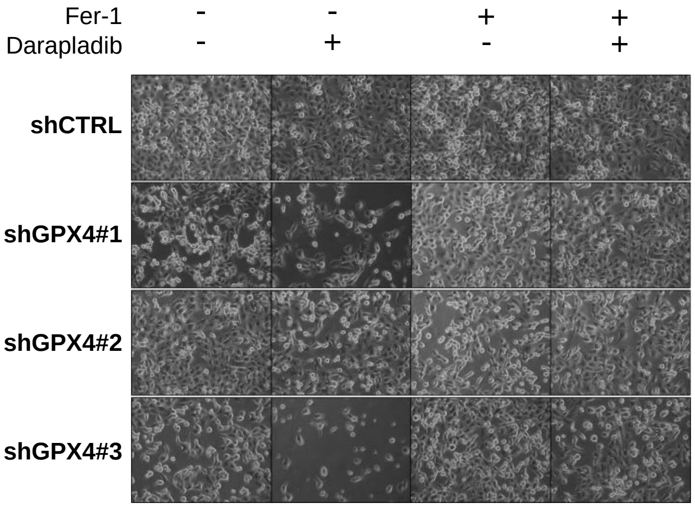


c

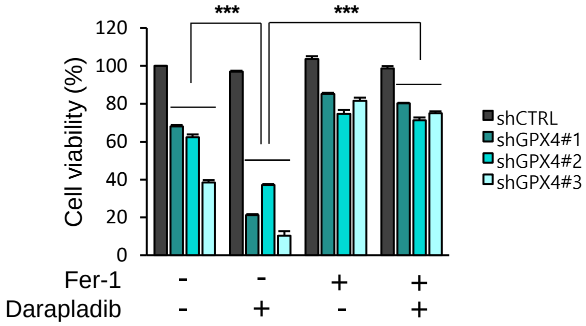

**Figure S13**

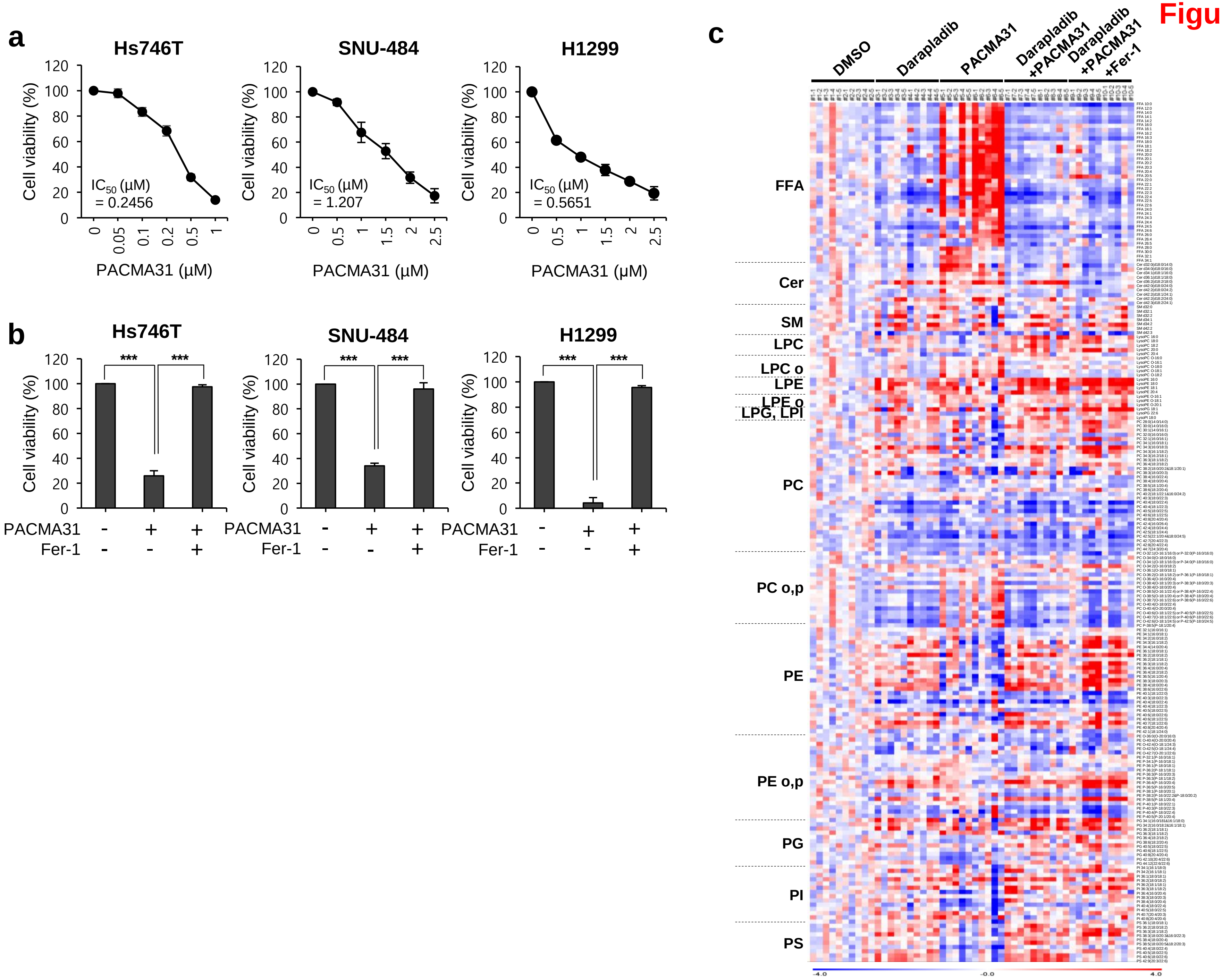
a
Hs746T
SNU-484
H1299
Cell viability (%)
IC50 (μM) = 0.2456
IC50 (μM) = 1.207
IC50 (μM) = 0.5651
PACMA31 (μM)
b
Hs746T
SNU-484
H1299
***
Cell viability (%)
PACMA31 - + +
Fer-1 - - +
c
DMSO
Darapladib
PACMA31
Darapladib +PACMA31
Darapladib +PACMA31 +Fer-1
FFA
Cer
SM
LPC
LPC o
LPE
LPE o
LPG, LPI
PC
PC o,p
PE
PE o,p
PG
PI
PS
-4.0
-0.0
4.0

### Figure S14

**a**

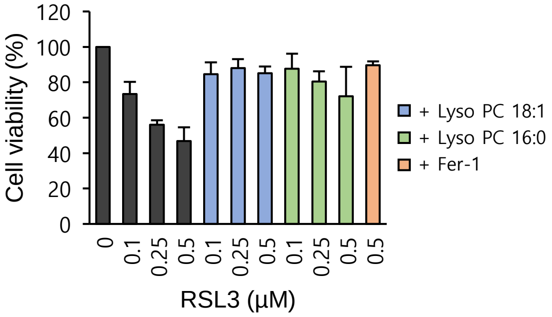

Cell viability (%)
120
100
80
60
40
20
0
0
0.1
0.25
0.5
0.1
0.25
0.5
0.1
0.25
0.5
0.5
RSL3 (μM)
+ Lyso PC 18:1
+ Lyso PC 16:0
+ Fer-1

### Figure S15

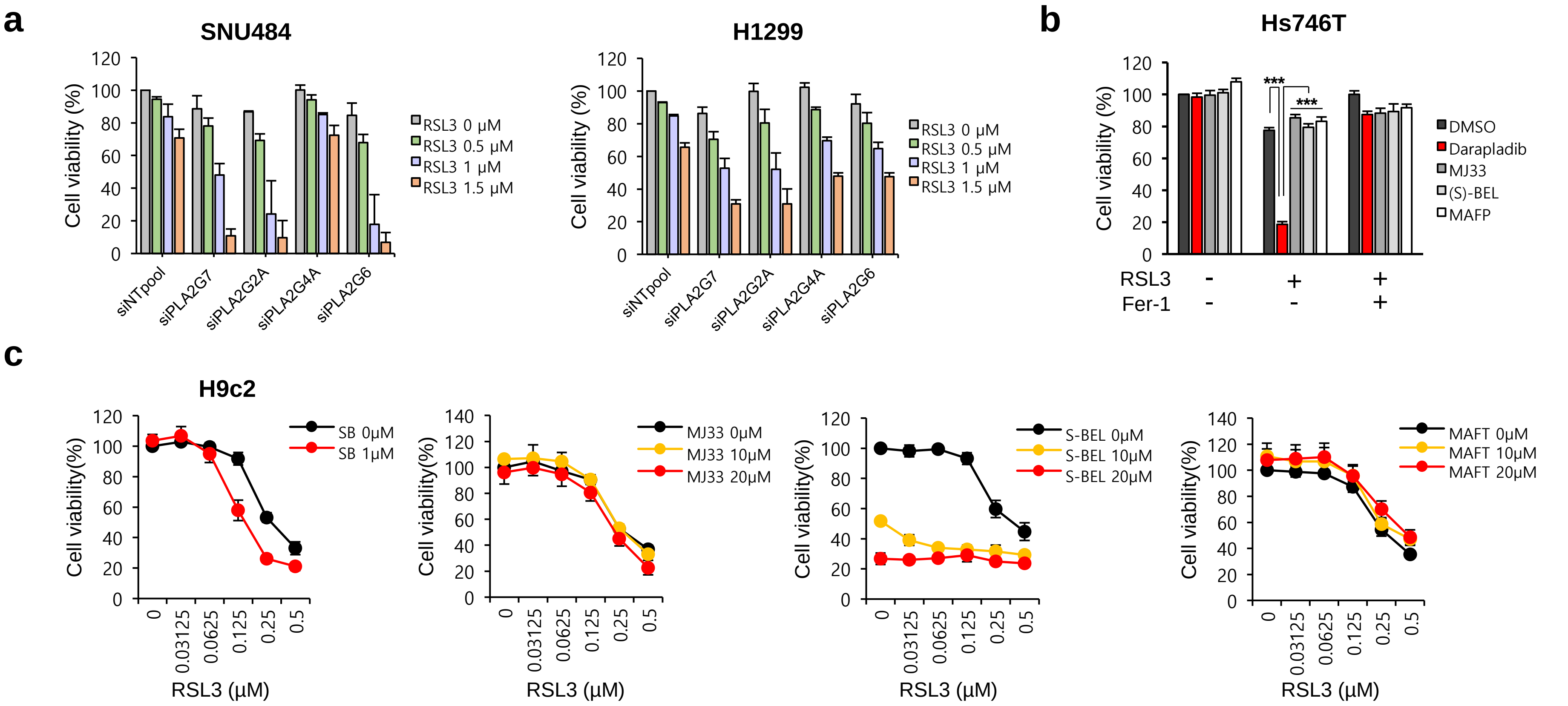

a
SNU484
H1299
Cell viability (%)
RSL3 0 μM
RSL3 0.5 μM
RSL3 1 μM
RSL3 1.5 μM
siNTpool
siPLA2G7
siPLA2G2A
siPLA2G4A
siPLA2G6
b
Hs746T
Cell viability (%)
DMSO
Darapladib
MJ33
(S)-BEL
MAFP
***
***
RSL3 - + +
Fer-1 - - +
c
H9c2
Cell viability(%)
SB 0μM
SB 1μM
MJ33 0μM
MJ33 10μM
MJ33 20μM
S-BEL 0μM
S-BEL 10μM
S-BEL 20μM
MAFT 0μM
MAFT 10μM
MAFT 20μM
0
0.03125
0.0625
0.125
0.25
0.5
RSL3 (μM)
